# Locus coeruleus MRI contrast is associated with cortical thickness in older adults

**DOI:** 10.1101/2020.03.14.991596

**Authors:** Shelby L. Bachman, Martin J. Dahl, Markus Werkle-Bergner, Sandra Düzel, Caroline Garcia Forlim, Ulman Lindenberger, Simone Kühn, Mara Mather

## Abstract

There is growing evidence that neuronal integrity of the noradrenergic locus coeruleus (LC) is important for later-life cognition. Less understood is how LC integrity relates to brain correlates of cognition, such as brain structure. Here, we examined the relationship between cortical thickness and a measure reflecting LC integrity in older (n = 229) and younger adults (n = 67). Using a magnetic resonance imaging sequence which yields high signal intensity in the LC, we assessed the contrast between signal intensity of the LC and that of neighboring pontine reference tissue. The Freesurfer software suite was used to quantify cortical thickness. LC contrast was positively related to cortical thickness in older adults, and this association was prominent in parietal, frontal, and occipital regions. Brain regions where LC contrast was related to cortical thickness include portions of the frontoparietal network which have been implicated in noradrenergically modulated cognitive functions. These findings provide novel evidence for a link between LC structure and cortical brain structure in later adulthood.

## 1. Introduction

A central goal of aging research is to identify factors which protect against age-related cognitive decline. Recently, the locus coeruleus-norepinephrine (LC-NE) system has been postulated as one such factor shaping later-life cognition (Mather & Harley, 2016; Robertson, 2013; Weinshenker, 2018). NE, an arousal-related neurotransmitter, facilitates neural and cognitive processes involved in vigilance, attention, and memory (Sara, 2009). The brain’s primary source of NE is the LC, a brainstem nucleus located adjacent to the lateral floor of the fourth ventricle (Sara & Bouret, 2012). The LC is densely packed with neurons which release NE throughout the brain (Schwarz & Luo, 2015); through these distributed projections, the LC stimulates neural responses which promote processing of relevant or salient information (Berridge & Waterhouse, 2003; Mather et al., 2016). Initial evidence that LC neuronal integrity relates to cognitive performance came from a longitudinal study which indicated that LC neuronal density – quantified through post-mortem brain autopsies – was higher in individuals who exhibited slower cognitive decline over a 6-year period before death (Wilson et al., 2013).

Studying the LC in vivo in humans has historically been challenging due to the small size of this nucleus. In recent years, however, the human LC has been quantified using magnetic resonance imaging (MRI) protocols, including turbo spin echo (TSE) and magnetization transfer sequences (Betts et al., 2019b; Liu et al., 2017; Sasaki et al., 2006), in which the LC is visible as a hyperintense region which is distinguishable from surrounding tissue (Keren et al., 2009; Shibata et al., 2006). Signal intensity contrast between the LC and surrounding pontine tissue in these sequences – henceforth referred to as LC contrast – has been validated through histological examination, with locations of high LC contrast corresponding to those of cells within the LC (Cassidy et al., 2019; Keren et al., 2015).

Studies using MRI to assess LC contrast have pointed to the importance of the LC in cognitive aging. In one study, LC contrast was positively related to cognitive reserve (Clewett et al., 2016), one indicator of the brain’s resilience against age-related pathology (Stern, 2009). LC contrast has also been linked to memory ability, with older adults with higher LC contrast having better episodic memory performance than those with lower contrast (Dahl et al., 2019; Hämmerer et al., 2018; Liu et al., 2020).

Additionally, individuals with Alzheimer’s disease (Betts et al., 2019a) and Parkinson’s disease (Liu et al., 2017) exhibited reduced LC contrast compared to healthy older adults. Together, these studies suggest a potentially neuroprotective role of LC integrity, as indexed by LC contrast, in older adulthood.

Of relevance to later-life cognition, NE released from the LC may protect against age-related neuropathology. For example, NE protects against neuroinflammation (Feinstein et al., 2016), regulates microglial functions which facilitate the clearance of amyloid beta (Heneka et al., 2010), contributes to increased expression of neurotrophic factors (Braun et al., 2014), reduces damage from neurotoxicity (Madrigal et al., 2007), and alleviates oxidative stress (Troadec et al., 2001). In addition, NE modulates a host of cognitive processes which typically decline with age, including selective attention (Mather et al., 2016; Sara, 2009), working and episodic memory (Berridge & Waterhouse, 2003; O’Dell et al., 2015), and cognitive flexibility (Lapiz & Morilak, 2006). Thus, for older adults, having relatively preserved neuronal integrity of the LC could lead to enhanced protection against brain pathology and enhanced noradrenergic modulation of cognition (Mather & Harley, 2016; Robertson, 2013).

If the LC has a neuroprotective role in older adulthood, LC integrity should be associated with cortical integrity. Older adults with greater cortical thickness have better fluid cognitive ability compared to those with lower cortical thickness (Fjell et al., 2006), and reductions in cortical thickness and volume explain a large amount of the variance in age-related cognitive decline (Fjell & Walhovd, 2010). To test the possibility that LC integrity is associated with brain structure, we examined the relationship between LC contrast quantified from TSE MRI scans and cortical thickness in older and younger adults. We predicted that LC contrast would be positively associated with cortical thickness and that this association would be strongest in brain regions which are recruited by NE modulated cognitive functions, including attention, cognitive flexibility, and memory (Corbetta et al., 2008; Robertson, 2014; Sara & Bouret, 2012).

## 2. Methods

### 2.1. Participants

Data were collected as part of the Berlin Aging Study II (BASE-II; Bertram et al., 2014; https://www.base2.mpg.de). A MR-eligible subsample of the BASE-II cohort for whom TSE scans were collected during the study’s second MR measurement-timepoint was selected for this study (n = 323). This subset included older and younger adults with no history of neurological disorder, psychiatric disorder, or head injury. All participants were right-handed with normal or corrected-to-normal vision. Data for eligible participants were acquired at the Max Planck Institute for Human Development in Berlin, Germany. The Ethics Committees of the German Psychological Society and of the Max Planck Institute for Human Development approved the MRI procedure. All participants provided written, informed consent prior to participation.

Participants were excluded if they did not have complete neural (n = 19) or demographic (n = 1) data. Following visual inspection of TSE scans, additional participants were excluded due to incorrect scan positioning (n = 3), motion artefact (n = 2), or incorrect placement of the LC search space (n = 1; see Section 2.3). One additional participant was excluded for having an inadequate cortical reconstruction. The final sample, described in Table 1, consisted of 229 older adults (82 females) and 67 younger adults (22 females). Older and younger adults did not differ significantly in terms of gender distribution, χ^2^(1, 296) = 0.092, *p* = .762, or mean years of education, *t*(104.4) = −0.347, *p* = .729. The older adult cohort had a significantly higher mean body mass index than did the younger adults, *t*(82.3) = 6.83, *p* < .001. Mini Mental State Examination (MMSE) scores for the older adult cohort ranged from 22 to 30 (mean = 28.6, SD = 1.30). No exclusions were performed based on MMSE scores. Excluding the two older adult participants who scored less than 25 on the MMSE did not change the pattern of results. The final sample exhibited a 99% overlap with the sample of BASE-II analyzed by Dahl et al. (2019).

**Table 1.** Sample characteristics

|  | Younger adults<br>(*n* = 67; 22 females) |  |  |  | Older adults<br>(*n* = 229; 82 females) |  |  |  |
| --- | --- | --- | --- | --- | --- | --- | --- | --- |
|  | Mean | SD | Min | Max | Mean | SD | Min | Max |
| Age | 32.6 | 3.62 | 25 | 40 | 72.3 | 4.10 | 63 | 83 |
| BMI | 23.0 | 3.75 | 11.6 | 31.9 | 26.7 | 3.49 | 19.1 | 37.9 |
| Education | 14.3 | 2.54 | 10 | 18 | 14.1 | 2.94 | 7 | 18 |
| MMSE | - | - | - | - | 28.6 | 1.30 | 22 | 30 |

*Note.* Age and education are expressed in years. BMI = body mass index; MMSE = Mini Mental State Examination; SD = standard deviation.
*2.2. MRI Data Acquisition*

### 2.2. MRI Data Acquisition

MRI data were acquired using a 3T Siemens Magnetom TIM Trio scanner with a 12-channel head coil. Only sequences used for the present analyses are described in this section. A three-dimensional, T1-weighted magnetization prepared rapid acquisition gradient-echo (MPRAGE) MRI sequence was applied in the sagittal plane (TR = 2500 ms, TE = 4.77 ms, TI = 1100 ms, scan flip angle = 7°; bandwidth = 140 Hz/pixel, field of view = 256 mm, number of slices = 192, isometric voxel size = 1 mm^3^, echo spacing = 10.9 ms; duration = 9:20 min).

Based on the MPRAGE sequence, a two-dimensional TSE sequence was applied by aligning the field of view orthogonally with respect to the anatomic axis of each participant’s brainstem (TR = 600 ms; TE = 11 ms; refocusing flip angle = 120°; no explicit MT saturation; bandwidth = 287 Hz/pixel; field of view = 256 mm; voxel size = 0.5 × 0.5 × 2.5 mm; echo spacing = 10.9 ms; duration = 2 × 5:54 min). This sequence included ten axial slices and a 20% gap between slices in the *z*-dimension, encompassing the entire pons. For some participants, fewer slices were acquired due to specific absorption rate limits being exceeded (for details, see Dahl et al., 2019; for a discussion, see Betts et al., 2019b). Four online averages were performed, yielding two TSE scans per participant. TSE scans from randomly selected younger and older adult participants are displayed in Figure S1.

### 2.3. LC Signal Intensity Assessment

We obtained LC contrast estimates using a semi-automated method, as described by Dahl et al. (2019). This approach yielded a LC location probability map which corresponded to published LC masks (Betts et al., 2017; Keren et al., 2009; Liu et al., 2019) as well as intensity estimates which corresponded to those determined through manual LC delineation. In this approach, Advanced Normalization Tools (Version 2.1; Avants et al., 2009) was used to generate a template brainstem volume by aligning and pooling TSE scans across participants (Figure 1A-C). The template was thresholded based on the signal intensity of a reference region in the neighboring dorsal pontine tegmentum (DPT), yielding a binarized LC search space (Figure 1D, 1E). After verifying that this search space encompassed the LC on a group level, this search space was applied as a mask on individual, template-coregistered TSE scans. The intensity and location of the maximum-intensity voxel within the masked region was extracted for each slice in the *z* (rostrocaudal, in whole-body coordinates) dimension of each masked volume. Likewise, a binarized mask of the DPT reference region was applied to template-coregistered TSE scans, and the intensity and location of the maximum-intensity DPT voxel was extracted for each slice. LC contrast values were calculated for each slice as a ratio (Betts et al., 2019b; Sasaki et al., 2006):

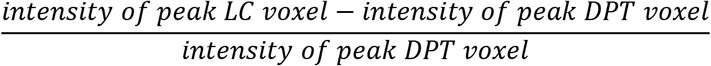

**Figure 1.**
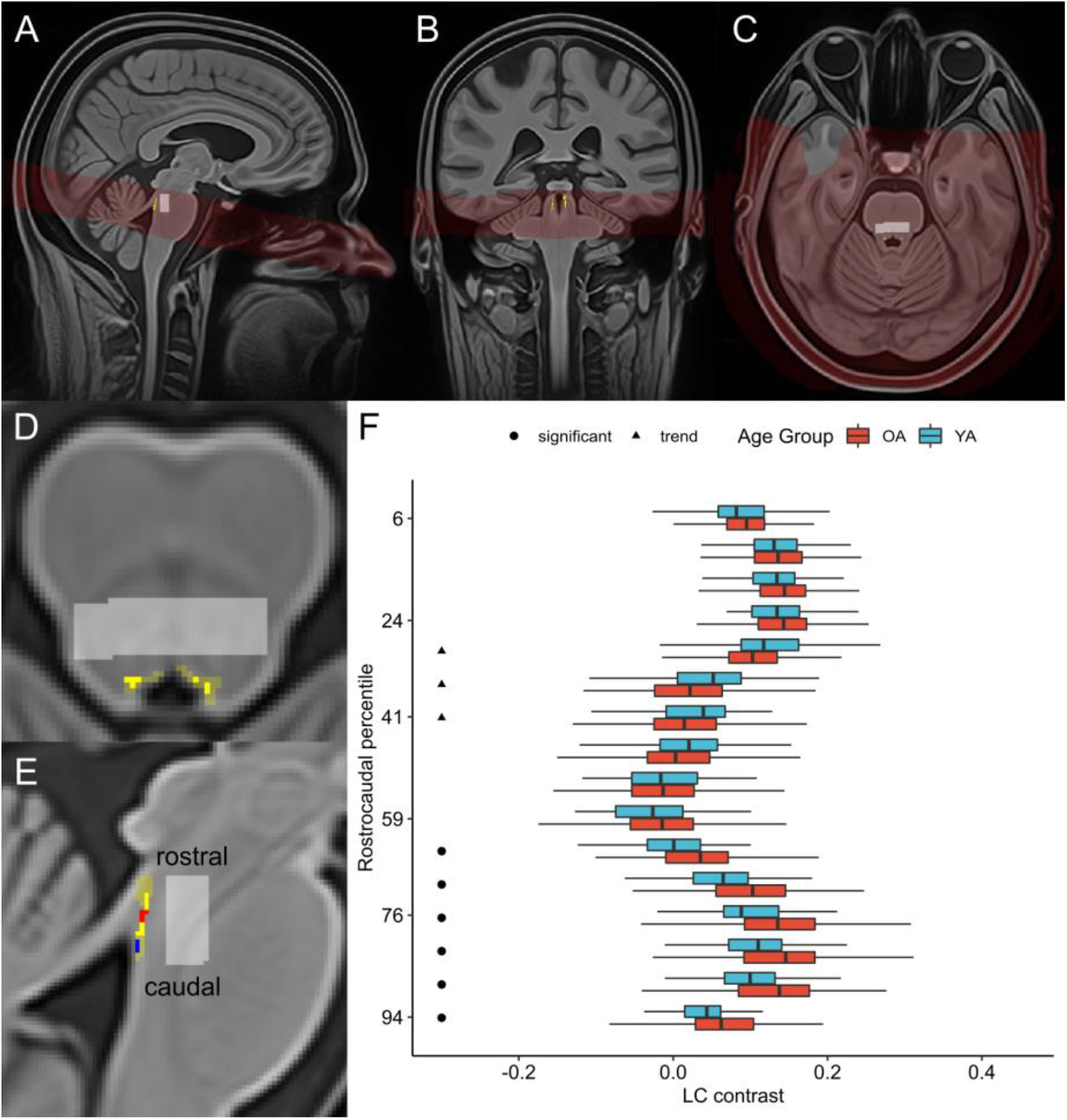
(A) Sagittal, (B) coronal, and (C) axial views of brainstem template volume (in red; generated from all participants’ turbo spin echo scans) overlaid onto whole-brain template (generated from all participants’ MPRAGE scans), with locus coeruleus (LC) probability map in common template space (Dahl et al., 2019) displayed in yellow. Pontine reference region in common template space is displayed in light gray. (D) Axial and (E) sagittal views of template brainstem volume. LC probability map (opaque yellow) is overlaid onto LC search space (translucent yellow), both of which are shown in common template space. Pontine reference region in common template space is displayed in light gray. In (E), rostral and caudal clusters within LC probability map are depicted in red and blue, respectively. (F) Boxplots depicting LC contrast values along the LC’s rostrocaudal extent are displayed for older adults (OA; red) and younger adults (YA; blue). The y-axis is arranged from most caudal (100) to most rostral (0) LC segment. For visual clarity, outliers – defined as data points falling outside the interval defined by 1.5 times the interquartile range (i.e. below the 25% quantile or above the 75% quantile) – are omitted. A cluster-wise permutation test was performed to identify clusters of slices where contrast differed significantly by age group. Circles indicate a caudal cluster of slices where older adults had significantly higher contrast than younger adults, and triangles indicate a rostral cluster of slices where there was a trend towards older adults having lower contrast than younger adults.

LC contrast values for each hemisphere were assessed in each participant’s two TSE scans separately, and contrast values were subsequently averaged within participants. Estimates of left and right LC contrast, reflecting the peak LC contrast value across all MRI dimensions in each hemisphere, were extracted. A structural equation model was used to estimate an overall value of LC contrast for each participant using left and right LC contrast as indicators (Dahl et al., 2019); the resulting estimates will henceforth be referred to as overall LC contrast values. In addition, for analyses of contrast along the LC’s rostrocaudal axis, we averaged LC contrast values across hemispheres within each slice to obtain slice-wise values of LC contrast for each participant.

### 2.4. Cortical Thickness Assessment

Cortical reconstruction for all T1-weighted anatomical images was performed with the Freesurfer software suite (Version 5.3.0; https://surfer.nmr.mgh.harvard.edu). Freesurfer’s automated processing pipeline has been described previously and entails motion correction, brain extraction, registration to Talairach coordinates, bias field estimation and correction, and intensity normalization (Dale et al., 1999; Fischl et al., 1999). These operations were applied, and pial and white matter surfaces were generated based on intensity gradients across tissue classes. One researcher (SB) blinded to participants’ age reviewed all resulting surfaces overlaid onto their respective anatomical images to identify segmentation inaccuracies which could bias subsequent thickness estimates (McCarthy et al., 2015). Only segmentation errors which did not resolve within 5 slices in any direction were manually corrected. A total of 122 (95 older adult, 27 younger adult) reconstructions contained inaccuracies on the pial surface and required that voxels be manually erased or included. The proportion of edited reconstructions did not differ by age group, χ^2^(1, 296) = 0.001, *p* = .974. For group analysis, all surfaces were tessellated and registered to a spherical atlas. Neuroanatomical labels corresponding to sulcal and gyral regions were automatically assigned to each vertex (Desikan et al., 2006; Fischl et al., 2004). At each vertex, cortical thickness was calculated as the shortest distance between the pial and white matter surfaces (Fischl & Dale, 2000). Thickness maps were smoothed with a circularly symmetric Gaussian kernel of 10 mm at full width half maximum, which was chosen to balance the increase in signal-to-noise afforded by smoothing with the risk of false alarms and loss of spatial precision which comes along with larger smoothing kernels. To probe relations between LC contrast and thickness of the entire cortical surface, a single, global thickness value was computed for each participant by averaging thickness over the left and right hemispheres and using the surface area of each hemisphere as a weighting factor.

### 2.5. Statistical Analysis

#### 2.5.1. LC contrast and global cortical thickness in the sample

Sample characteristics of younger and older adults were examined using independent samples *t*-tests with equal variances not assumed, chi-square independence tests, and Pearson correlation analyses. Freesurfer’s group analysis stream was used to examine cortical regions in which thickness differed between older and younger adults. Previous studies have reported age-related differences in LC contrast along the LC’s rostrocaudal extent (Betts et al., 2017; Dahl et al., 2019; Liu et al., 2019; Manaye et al., 1995). To confirm that the slight sample difference in this study did not change the topographic results reported in Dahl et al. (2019), we reimplemented the nonparametric, cluster-wise permutation test to identify clusters of slices where LC contrast differed significantly between younger and older adults (Maris & Oostenveld, 2007). This approach is described in detail in the Supplementary Methods (Section 1.2). To test for reliable age differences in LC contrast according to topography, we then performed a mixed analysis of variance with age group (younger, older) and topography (one level per identified cluster) as factors.

#### 2.5.2. Analysis of associations between LC contrast and global cortical thickness

We used multiple linear regression to determine whether LC contrast was associated with global cortical thickness and whether this association depended on age. Specifically, we constructed a regression model with overall LC contrast and chronological age as predictors of global thickness. In addition, to determine whether the effect of LC contrast on global thickness differed between older and younger adults, we included the interaction effect of overall LC contrast and age group (coded as 1=older adults, −1=younger adults) as a predictor. Multiple linear regression analyses were subsequently conducted to examine the effects of overall LC contrast and age on global thickness in each age group separately. Each predictor was standardized by mean centering and dividing by its standard deviation; thus, results of these analyses are reported as standardized regression coefficients and standard error values. For analyses involving both age groups, predictors were standardized based on values from the entire sample; for analyses of each age group separately, predictors were standardized for each group separately. To determine whether associations between LC contrast and thickness differed based on LC topography, we repeated these analyses for each set of cluster-specific LC contrast values. We also performed supplementary analyses for left and right LC contrast to examine potential effects of LC laterality. In addition, exploratory analyses indicated significant sex differences in overall LC contrast and global cortical thickness, so we performed separate regression analyses accounting for sex and its interaction effects with other predictors on thickness (Supplementary Methods, Section 1.3). All descriptive statistics and regression analyses were performed in R (Version 3.6.2; R Core Team, 2019).

#### 2.5.3. Vertex-wise analysis of associations between LC contrast and cortical thickness

To examine locations on the cortical surface where LC contrast was associated with thickness, we fit a series of vertex-wise general linear models using Freesurfer’s group analysis stream. First, we tested whether there were cortical regions where the association between overall LC contrast and thickness differed in older and younger adults, while regressing out the effect of chronological age. Then, for each age group separately, we modeled cortical thickness at each vertex as a function of overall LC contrast while regressing out the effect of chronological age. To determine whether associations with thickness depended on LC laterality, we performed supplementary vertex-wise analyses for left and right LC contrast separately. In addition, to determine whether regionally specific associations between cortical thickness and LC contrast differed based on LC topography, we performed these vertex-wise analyses again using each set of cluster-specific LC contrast values. Finally, in line with analyses of global thickness, we conducted supplementary vertex-wise analyses accounting for sex (Supplementary Methods, Section 1.4). For all vertex-wise analyses, significance maps were thresholded at vertex-wise *p* < 0.05, and cluster-wise correction for multiple comparisons was performed using a Monte Carlo Null-Z simulation with 10000 iterations (Hagler et al., 2006). This entailed generating a distribution of the maximum cluster size under the null hypothesis, which was used to compute cluster-wise *p*-values. A cluster-wise threshold of *p* < 0.05 was applied to include only those clusters unlikely to appear by chance.

## 3. Results

### 3.1. LC contrast & global cortical thickness in the sample

Overall LC contrast was comparable in younger and older adults, *t*(88.6) = −1.24, *p* = .218, and was not significantly correlated with chronological age in either age group (Figure S2). Across the sample, contrast was higher for the left compared to the right LC, *t*(295) = 6.72,*p* < .001 (Figure S3). Consistent with the findings of Dahl et al. (2019), we detected age group differences in LC contrast along the LC’s rostrocaudal extent (Figure 1F). Specifically, we detected a cluster of 6 caudal slices where older adults had greater LC contrast than younger adults, *p* = .002, as well as a cluster of 3 rostral slices where there was a trend towards older adults having lower LC contrast than younger adults, *p* = .083. A subsequent analysis of variance indicated a significant interaction effect of age group and topography (rostral/caudal) on LC contrast, *F*(1,294) = 25.1,*p* < .001. As anticipated, global cortical thickness was significantly lower in older adults compared to younger adults, *t*(108.9) = −9.61, *p* < .001, and was negatively correlated with age in both age groups (Figure S4A). Thickness was lower in older relative to younger adults in many cortical regions (Figure S4B). Exploratory analyses indicated significant sex differences in both overall LC contrast and global cortical thickness (Figure S5).

### 3.2. Analysis of the association between LC contrast and global cortical thickness

Figure 2 depicts associations between LC contrast and cortical thickness in older and younger adults. Multiple linear regression analysis indicated a significant interaction effect between overall LC contrast and age group on global thickness, *β* = 0.138, *p* = .009, indicating that the association between overall LC contrast and global thickness differed significantly in older and younger adults (Table 2A). In older adults, higher overall LC contrast, *β* = 0.216, *p* < .001, and lower age, *β* = −0.153, *p* = .018, were significantly associated with greater global thickness (Table 2B). In younger adults, neither overall LC contrast nor age was significantly associated with global thickness (Table 2C). Exploratory analyses indicated that in older adults, global thickness was positively related to both left and right LC contrast (Supplementary Results, Section 2.3.1).

**Figure 2.**
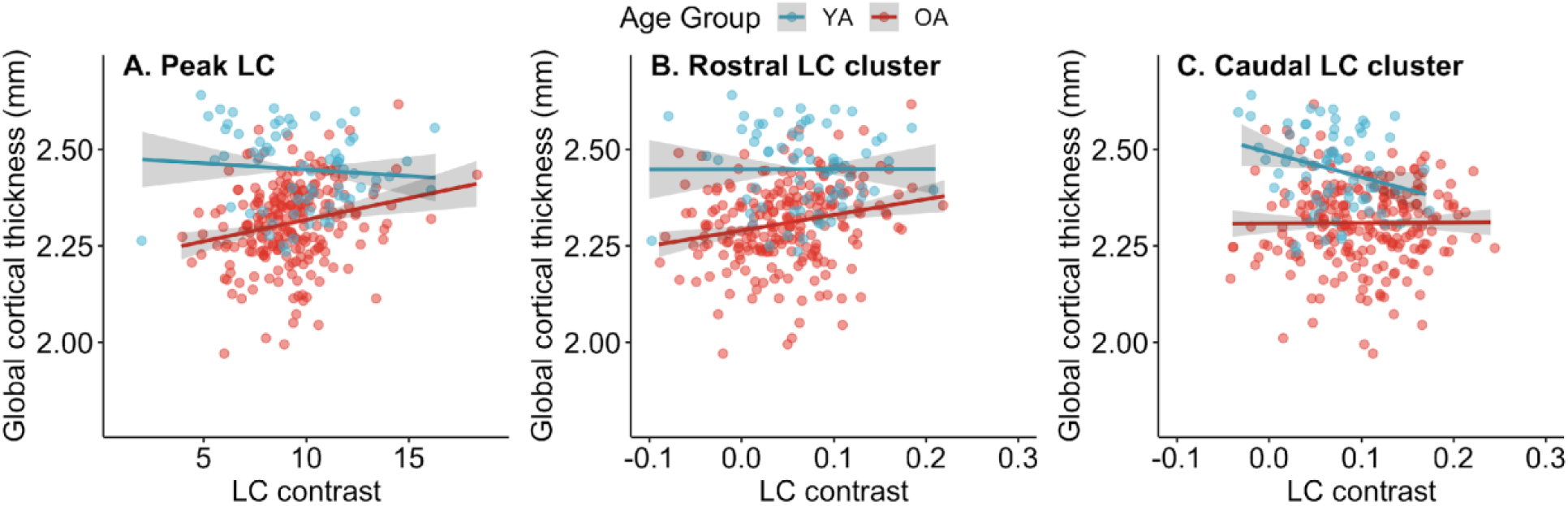
Associations between locus coeruleus (LC) contrast and global cortical thickness. Associations with global cortical thickness are displayed separately for older adults (OA) and younger adults (YA) for (A) overall LC contrast (reflecting contrast of peak LC voxels across x-, y-, and z-dimensions), (B) contrast of the rostral LC, and (C) contrast of the caudal LC. Points indicate raw data. Model fit lines reflect the association between LC contrast and global cortical thickness after regressing out the effect of chronological age on thickness. 95% confidence intervals are shown in gray. Fit lines and confidence intervals were calculated using the mean value of chronological age within each age group. Differences in scales on the x-axes are due to overall LC being calculated as a latent variable in a structural equation modeling framework, whereas rostral and caudal LC contrast were calculated by averaging signal intensity contrast ratios.

**Table 2.** Results of multiple linear regression analyses examining the relationship between overall locus coeruleus contrast and global cortical thickness

| Predictor | $\beta$ | SE | 95% CI | $t$ | $p$ |
| --- | --- | --- | --- | --- | --- |
| A. Full sample, $F(3, 292)=39.38, p<.001, R^2=0.288$, adjusted $R^2 = 0.281$ | | | | | |
| Overall LC contrast | 0.069 | 0.052 | -0.033, 0.172 | 1.33 | .185 |
| **Age** | -0.506 | 0.050 | -0.604, -0.408 | -10.2 | <.001 |
| **Overall LC contrast \* Age Group** | 0.138 | 0.052 | 0.035, 0.241 | 2.64 | .009 |
| B. Older adults, $F(2, 226)=9.14, p<.001, R^2=0.075$, adjusted $R^2=0.067$ | | | | | |
| **Overall LC contrast** | 0.216 | 0.064 | 0.090, 0.342 | 3.37 | <.001 |
| **Age** | -0.153 | 0.064 | -0.280, -0.027 | -2.39 | .018 |
| C. Younger adults, $F(2, 64)=1.72, p=.187, R^2=0.051$, adjusted $R^2=0.021$ | | | | | |
| Overall LC contrast | -0.087 | 0.123 | -0.332, 0.158 | -0.71 | .482 |
| Age | -0.198 | 0.123 | -0.443, 0.047 | -1.62 | .111 |

*Note.* Regression coefficients and standard errors are standardized values. Age group was coded as: older adults = 1, younger adults = -1. CI = confidence interval; LC = locus coeruleus; SE = standard error.

To examine effects of LC topography on the association between LC contrast and thickness, we performed analogous regression analyses using rostral and caudal values of LC contrast, respectively, as predictors instead of overall LC contrast. An analysis of the full sample indicated a trend towards an interaction effect between rostral LC contrast and age group, *β* = 0.103, *p* = .082 (Table 3A). Although this result did not indicate a significant difference in the rostral LC contrast-global thickness association by age group, for consistency with analyses of overall LC contrast, we performed subsequent regression analyses in each age group separately. These analyses demonstrated that the rostral LC-thickness association was driven by older adults, with rostral LC contrast demonstrating a significantly positive association with global thickness in older adults, *β* = 0.214,*p* < .001 (Table 3B), but not in younger adults, *β* = 0.002, *p* = .990 (Table 3C).

**Table 3.** Results of multiple linear regression analyses examining the relationship between rostral locus coeruleus contrast and global cortical thickness

| Predictor | $\beta$ | SE | 95% CI | $t$ | $p$ |
| --- | --- | --- | --- | --- | --- |
| A. Full sample, $F(3, 292)=38.98$, $p<.001$, $R^2=0.286$, adjusted $R^2=0.279$ | | | | | |
| Rostral LC contrast | 0.088 | 0.060 | -0.029, 0.204 | 1.48 | .139 |
| **Age** | -0.503 | 0.051 | -0.602, -0.403 | -9.93 | <.001 |
| Rostral LC contrast * Age Group | 0.103 | 0.059 | -0.013, 0.220 | 1.75 | .082 |
| B. Older adults, $F(2, 226)=9.08$, $p<.001$, $R^2=0.074$, adjusted $R^2=0.066$ | | | | | |
| **Rostral LC contrast** | 0.214 | 0.064 | 0.088, 0.341 | 3.35 | <.001 |
| **Age** | -0.163 | 0.064 | -0.289, -0.036 | -2.54 | .012 |
| C. Younger adults, $F(2, 64)=1.46$, $p=.240$, $R^2=0.044$, adjusted $R^2=0.014$ | | | | | |
| Rostral LC contrast | 0.002 | 0.124 | -0.246, 0.250 | 0.01 | .990 |
| Age | -0.209 | 0.124 | -0.457, 0.039 | -1.69 | .097 |

*Note.* Regression coefficients and standard errors are standardized values. Age group was coded as: older adults = 1, younger adults = -1. CI = confidence interval; LC = locus coeruleus; SE = standard error.

A regression analysis examining the association between caudal LC contrast and global cortical thickness indicated a trend towards caudal LC contrast being negatively associated with global thickness, *β* = −0.128, *p* = .071, as well as a marginally significant interaction effect between caudal LC contrast and age group on global thickness, *β* = 0.135, *p* = .057 (Table 4A). In older adults, caudal LC contrast was not significantly associated with global cortical thickness, *β* = 0.006, *p* = .922 (Table 4B). However, in younger adults, higher caudal LC contrast was associated with lower global cortical thickness, *β* = −0.275, *p* = .023 (Table 4C). Results of analyses examining associations between LC contrast and global thickness accounting for sex are included in the Supplementary Results (Section 2.4.1).

**Table 4.**
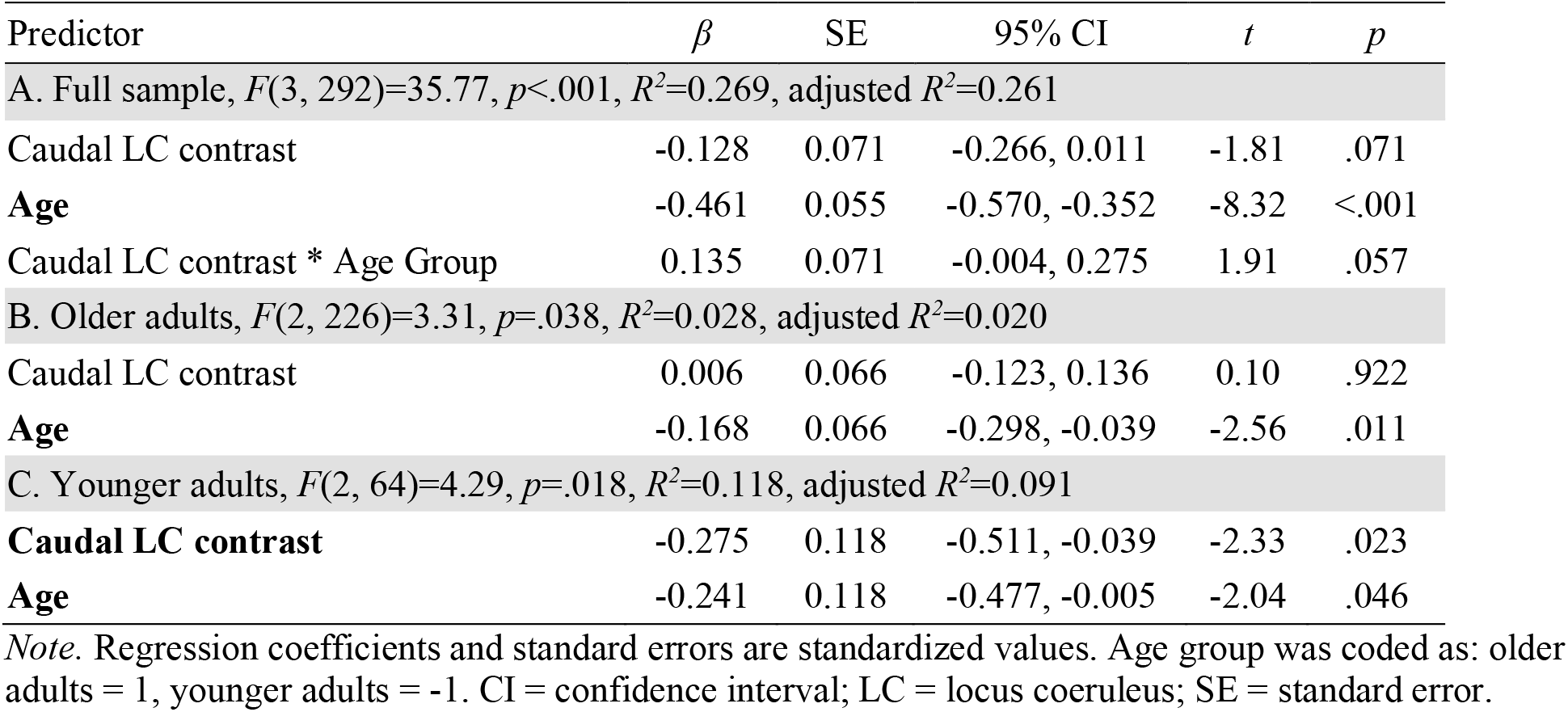
Results of multiple linear regression analyses examining the relationship between caudal LC contrast and global cortical thickness

### 3.3. Analysis of the association between LC contrast and regional cortical thickness

Using vertex-wise analyses, we detected ten cortical clusters where the association between overall LC contrast and thickness was more positive in older than younger adults; cluster details are presented in Table 5A. In an analysis of older adults only, we found a significant association between overall LC contrast and cortical thickness in fourteen clusters (Table 5B); in each of these clusters, overall LC contrast was positively associated with thickness. As shown in Figure 3, clusters identified in both analyses included regions in frontal, parietal, occipital, and temporal cortices. In an analysis of younger adults only, there were no clusters surviving multiple comparison correction in which overall LC contrast was associated with thickness. Uncorrected (vertex-wise *p* < .05) significance maps from analyses of overall LC contrast are displayed in Figure S6. Exploratory analyses indicated more widespread associations with thickness for left versus right LC contrast (Supplementary Results, Section 2.3.2).

**Figure 3.**
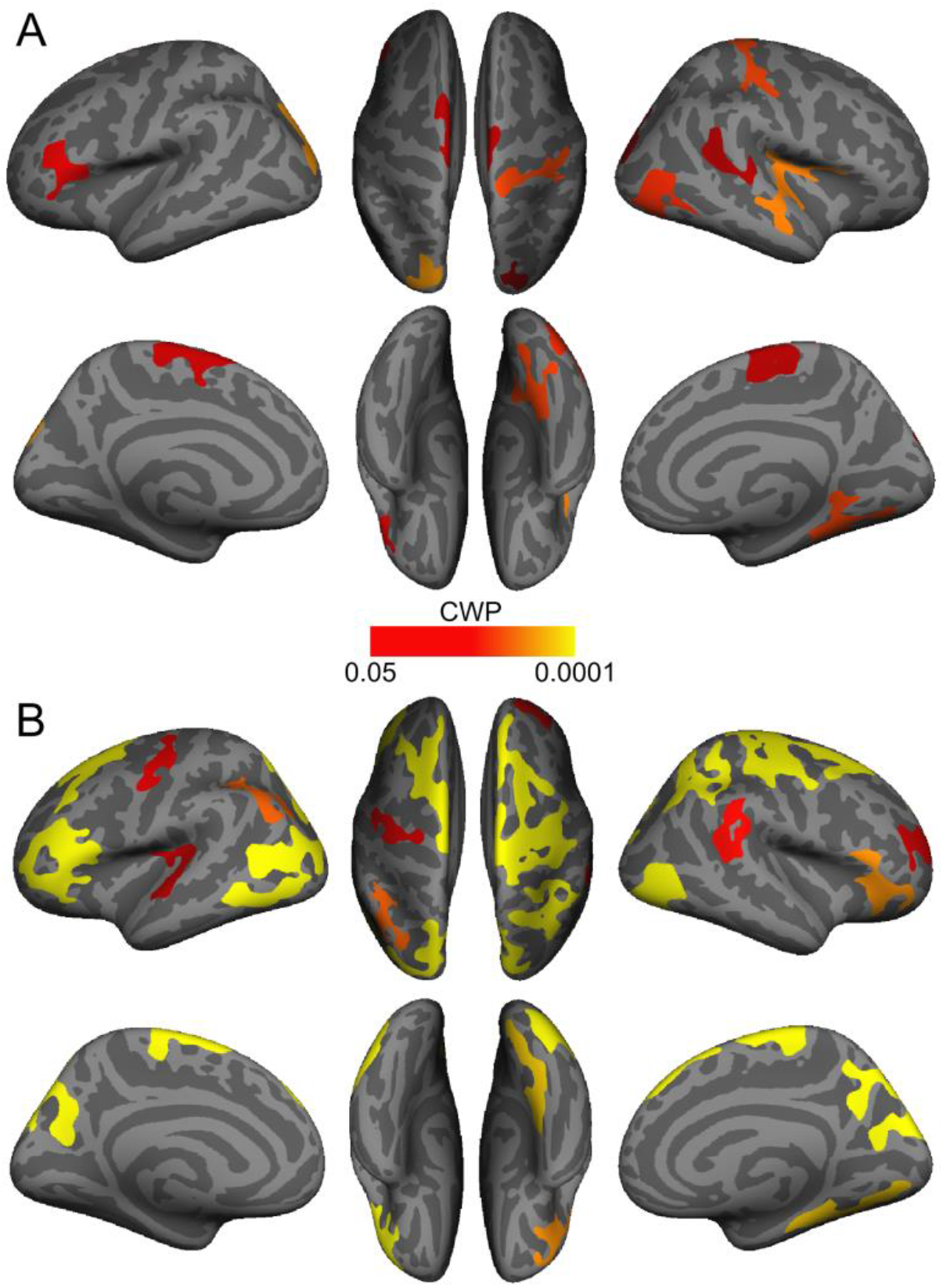
Cortical regions where the association between overall locus coeruleus contrast and thickness was more positive in older than in younger adults (A) and positive in older adults (B). Only clusters that survived cluster-wise correction for multiple comparisons (cluster-wise probability (CWP) < .05) are shown.

**Table 5.** Cortical clusters where the association between overall LC contrast and thickness was more positive in older than in younger adults (A) and positive in older adults (B)

| Region | Hemi-<br>sphere | Size<br>(mm<sup>2</sup>) | Peak vertex* |  |  | CWP** |
| --- | --- | --- | --- | --- | --- | --- |
|  |  |  | X | Y | Z |  |
| A. Overall LC - thickness association more positive in OA vs. YA |  |  |  |  |  |  |
| Superior parietal | Left | 1377.59 | -16.0 | -89.4 | 21.9 | 0.0005 |
| Rostral middle frontal | Left | 1028.72 | -35.5 | 31.4 | 14.8 | 0.0106 |
| Paracentral | Left | 953.99 | -8.1 | -22.4 | 65.1 | 0.0184 |
| Middle temporal | Right | 1447.35 | 51.1 | -12.5 | -20.7 | 0.0008 |
| Postcentral | Right | 1301.42 | 14.4 | -33.8 | 64.5 | 0.0021 |
| Fusiform | Right | 1282.95 | 36.0 | -43.5 | -13.9 | 0.0023 |
| Lateral occipital | Right | 1230.74 | 48.2 | -75.0 | 5.9 | 0.0032 |
| Paracentral | Right | 905.96 | 5.9 | -12.1 | 52.7 | 0.0251 |
| Banks superior temporal sulcus | Right | 903.84 | 46.0 | -42.9 | 13.9 | 0.0259 |
| Lateral occipital | Right | 811.95 | 23.9 | -90.1 | 18.9 | 0.0475 |
| B. Overall LC positively associated with thickness in OA |  |  |  |  |  |  |
| Inferior parietal | Left | 5334.79 | -44.4 | -70.2 | 9.4 | 0.0001 |
| Pars triangularis | Left | 2829.49 | -47.9 | 34.8 | -8.5 | 0.0001 |
| Superior frontal | Left | 2590.56 | -8.9 | 2.1 | 67.1 | 0.0001 |
| Inferior parietal | Left | 1350.49 | -43.3 | -63.9 | 38.9 | 0.0013 |
| Precentral | Left | 1039.94 | -34.7 | -21.0 | 49.8 | 0.0102 |
| Superior temporal | Left | 910.53 | -59.4 | -12.0 | -3.3 | 0.0244 |
| Precentral | Right | 5682.29 | 40.8 | -10.3 | 58.2 | 0.0001 |
| Precuneus | Right | 2556.13 | 6.7 | -57.4 | 32.3 | 0.0001 |
| Inferior parietal | Right | 2260.05 | 37.0 | -53.9 | 38.1 | 0.0001 |
| Lateral occipital | Right | 1783.10 | 40.8 | -85.3 | -13.0 | 0.0001 |
| Lingual | Right | 1602.38 | 20.4 | -74.2 | -8.8 | 0.0002 |
| Pars orbitalis | Right | 1389.28 | 45.8 | 30.2 | -12.3 | 0.0006 |
| Bank superior temporal sulcus | Right | 1154.07 | 48.8 | -40.6 | 12.5 | 0.0056 |
| Rostral middle frontal | Right | 1007.60 | 31.0 | 41.6 | 20.2 | 0.0138 |

*Note.* LC = locus coeruleus; OA = older adults; YA = younger adults. There were no clusters where overall LC contrast was associated with thickness in younger adults.
\*Talairach coordinates of vertex with peak CWP value.
\*\*Cluster-wise probability (CWP) resulting from cluster-wise Monte Carlo correction for multiple comparisons, reflecting the probability of the cluster appearing by chance. Only clusters with $CWP < .05$ are listed.

Analogous vertex-wise analyses indicated that, after correction for multiple comparisons, the association between rostral LC contrast differed in older and younger adults in one cluster in left superior parietal cortex (Table 6A); in this cluster, thickness was more positively associated with rostral LC contrast in older adults. In an analysis of older adults only, rostral LC contrast was positively associated with thickness in fifteen cortical clusters (Table 6B). Clusters where rostral LC contrast was associated with thickness in older adults, which are displayed in Figure 4A, included portions of parietal, frontal, and occipital cortices, and many overlapped with clusters where overall LC contrast was associated with thickness in older adults. In younger adults, there were no cortical clusters in which rostral LC contrast was significantly associated with thickness. Next, we found that the association between caudal LC contrast and thickness was more positive in older than younger adults in four cortical clusters (Table 6C). Analysis of older adults separately indicated that caudal LC contrast was only associated with thickness in one cluster in rostral middle frontal cortex (Table 6D), and in this cluster, caudal LC contrast was negatively associated with thickness (Figure 4B). Furthermore, in younger adults, caudal LC contrast was negatively associated with thickness in three clusters, which included regions in parietal and occipital cortices (Table 6E). Figures S7 and S8 depict uncorrected (vertex-wise *p* < .05) significance maps from analyses of rostral and caudal LC contrast, respectively, before cluster-wise correction for multiple comparisons. Results of supplementary vertex-wise analyses examining associations between LC contrast and thickness accounting for sex are included in the Supplementary Results (Section 2.4.2).

**Figure 4.**
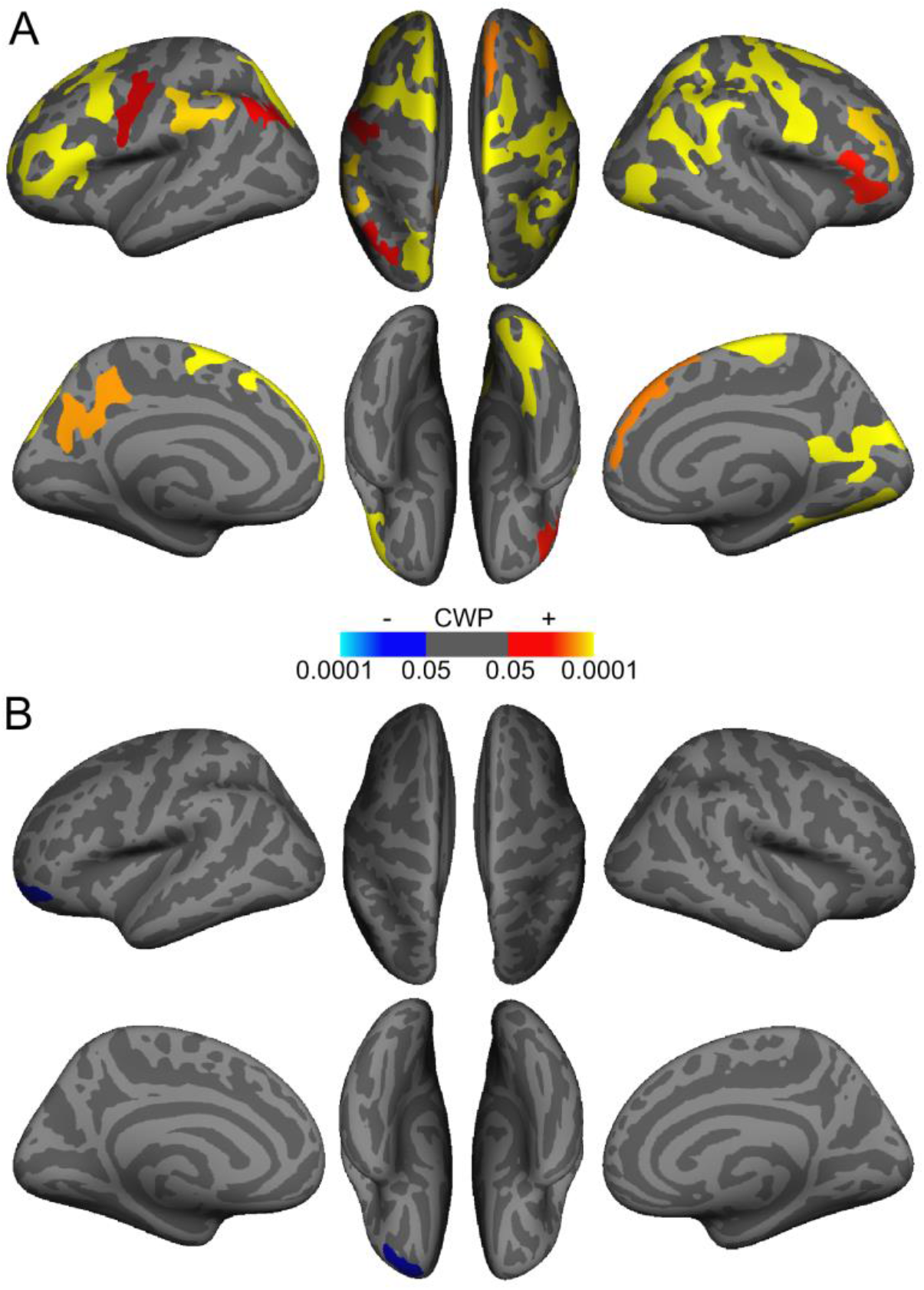
Cortical regions demonstrating an association between rostral (A) and caudal (B) locus coeruleus (LC) contrast and thickness in older adults. Warmer colors indicate clusters with a positive association between LC contrast and cortical thickness, and cooler colors indicate clusters with a negative association. Only clusters that survived cluster-wise correction for multiple comparisons (cluster-wise probability (CWP) <.05) are shown.

**Table 6.** Cortical clusters where the association between rostral LC contrast and thickness was more positive in older than in younger adults (A) and positive in older adults (B) and where the association between caudal LC contrast and thickness was more positive in older than in younger adults (C) and negative in older adults (D) and younger adults (E)

| Region | Hemi-<br>sphere | Association with<br>thickness | Size<br>(mm<sup>2</sup>) | Peak vertex* |  |  | CWP |
| --- | --- | --- | --- | --- | --- | --- | --- |
|  |  |  |  | X | Y | Z |  |
| A. Rostral LC contrast - thickness association more positive in OA vs. YA |  |  |  |  |  |  |  |
| Superior parietal | Left | OA > YA | 842.14 | -22.9 | -74.4 | 29.9 | 0.0387 |
| B. Rostral LC contrast associated with thickness in OA |  |  |  |  |  |  |  |
| Pars triangularis | Left | Positive | 2633.77 | -48.1 | 35.4 | -6.3 | 0.0001 |
| Caudal middle frontal | Left | Positive | 2288.88 | -39.6 | 11.4 | 51.4 | 0.0001 |
| Superior frontal | Left | Positive | 1958.83 | -10.1 | 39.7 | 47.9 | 0.0001 |
| Superior parietal | Left | Positive | 1811.48 | -10.2 | -87.1 | 29.1 | 0.0001 |
| Postcentral | Left | Positive | 1472.10 | -47.0 | -25.4 | 33.9 | 0.0002 |
| Precuneus | Left | Positive | 1391.17 | -8.2 | -45.3 | 44.3 | 0.0005 |
| Inferior parietal | Left | Positive | 1091.19 | -43.3 | -64.6 | 40.3 | 0.0070 |
| Precentral | Left | Positive | 974.53 | -45.2 | -10.9 | 39.7 | 0.0166 |
| Precentral | Right | Positive | 4717.72 | 41.0 | -9.8 | 58.5 | 0.0001 |
| Supramarginal | Right | Positive | 3809.56 | 57.9 | -39.9 | 35.6 | 0.0001 |
| Lateral occipital | Right | Positive | 2799.21 | 47.8 | -77.2 | 5.1 | 0.0001 |
| Isthmus cingulate | Right | Positive | 2070.17 | 8.9 | -53.3 | 9.6 | 0.0001 |
| Rostral middle frontal | Right | Positive | 1424.36 | 42.7 | 33.6 | 28.2 | 0.0002 |
| Superior frontal | Right | Positive | 1338.58 | 6.8 | 40.8 | 46.9 | 0.0008 |
| Pars triangularis | Right | Positive | 1096.91 | 54.2 | 26.6 | 8.1 | 0.0047 |
| C. Caudal LC contrast - thickness association more positive in OA vs. YA |  |  |  |  |  |  |  |
| Superior parietal | Left | OA > YA | 3135.14 | -19.1 | -86.0 | 18.8 | 0.0001 |
| Supramarginal | Right | OA > YA | 1278.17 | 45.1 | -28.6 | 37.7 | 0.0014 |
| Superior parietal | Right | OA > YA | 928.04 | 25.8 | -79.4 | 21.4 | 0.0227 |
| Paracentral | Right | OA > YA | 841.08 | 15.6 | -38.4 | 53.3 | 0.0409 |
| D. Caudal LC contrast associated with thickness in OA |  |  |  |  |  |  |  |
| Rostral middle frontal | Left | Negative | 857.23 | -19.1 | 57.7 | -11.9 | 0.0351 |
| E. Caudal LC contrast associated with thickness in YA |  |  |  |  |  |  |  |
| Lateral occipital | Left | Negative | 2959.26 | -34.0 | -87.6 | 9.6 | 0.0001 |
| Paracentral | Left | Negative | 1075.54 | -5.4 | -18.8 | 52.4 | 0.0065 |
| Superior parietal | Right | Negative | 2419.58 | 20.6 | -41.3 | 58.2 | 0.0001 |

*Note.* LC = locus coeruleus; OA = older adults; YA = younger adults. There were no clusters in younger adults where rostral LC contrast was associated with thickness.
\*Talairach coordinates of vertex with peak CWP value.
\*\*Cluster-wise probability (CWP) resulting from cluster-wise Monte Carlo correction for multiple comparisons, reflecting the probability of the cluster appearing by chance. Only clusters with CWP < .05 are listed.

## 4. Discussion

A growing body of literature suggests that the LC plays a role in cognition in later life, with recent studies linking LC MRI contrast (as an in vivo estimate of LC neuronal integrity) to cognitive outcomes in older adults (Clewett et al., 2016; Dahl et al., 2019; Hämmerer et al., 2018; Liu et al., 2020). In this study, we examined whether LC contrast was associated with cortical thickness, one aspect of brain structure and a documented brain indicator of cognitive ability in older adulthood (Fjell et al., 2006; Lee et al., 2016). We found that higher LC contrast was associated with greater average cortical thickness in older adults, even after regressing out effects of age on thickness. When we examined where this association was most evident in older adults, we found widespread regions in parietal, frontal, and occipital cortices in which higher LC contrast was associated with greater thickness. Together, these findings constitute novel evidence for a link between LC contrast and brain structure.

What biological processes might the relationship between LC contrast and cortical thickness reflect? One possibility is that having greater MR-indexed LC integrity could lead to higher levels of NE release, allowing the beneficial effects of *β*-adrenergic receptor activation to be realized throughout the brain and in turn to the preservation of brain structure. Many of the brain regions where LC contrast was associated with thickness belong to sensory, motor, association, and prefrontal cortices, all of which are innervated by the LC (Bouret & Sara, 2002; Chandler et al., 2014; Hirschberg et al., 2017). Critically, a number of regions where thickness was associated with LC contrast are contained within the brain’s frontoparietal network (FPN), which maintains flexible representations of environmental priority and internally- and externally-guided goals (Ptak, 2012; Zanto & Gazzaley, 2013). Frontoparietal regions show strong resting-state functional connectivity with the LC (Jacobs et al., 2018), and during arousal and attention, phasic LC activity can promote neural gain, modulating signal-to-noise ratio of neuronal activity within the FPN to direct processing toward salient and/or task-relevant stimuli (Aston-Jones & Cohen, 2005; Bouret & Sara, 2005; Corbetta et al., 2008; Lee et al., 2018). Thus, reduced or diminished LC-NE activity later in life, resulting from reduced LC neuronal integrity, could contribute to impairments in processes such as selective attention and working and episodic memory (Corbetta et al., 2008). Reduced noradrenergic signaling within brain regions implicated in NE-mediated functions, such as the FPN but also including occipital regions, could reduce NE’s protective effects in these brain regions, eventually permitting gray matter atrophy. On the other hand, preserved LC integrity could maintain the NE available to act on the FPN and other brain regions and, concurrently, more of NE’s protective effects being realized throughout the cortex.

We also found that the association between LC contrast and thickness depended on LC topography, as contrast of the rostral LC – but not the caudal LC – demonstrated associations with thickness in older adulthood. Previous work has demonstrated that cortical and thalamic projections from the LC – particularly those to frontal, sensory, and occipital cortices – tend to arise in its more rostral portion (Schwarz & Luo, 2015; Waterhouse et al., 1983). Likewise, projections from the LC to the basal forebrain tend to originate more from rostral than from caudal LC (España & Berridge, 2006). Supporting this idea, Dahl et al., (2019) found that rostral, but not caudal, LC contrast was associated with memory performance in older adults in a largely overlapping sample from BASE-II. Furthermore, consistent with findings of Liu et al. (2019), we found that rostral but not caudal LC exhibited age-related decline in contrast. On the other hand, contrast of the caudal LC demonstrated markedly different associations with thickness compared to contrast of the rostral LC. Whereas in older adults caudal LC contrast was largely unrelated to thickness, it was negatively associated with thickness in younger adults. As previous studies have not reported associations between cognition and caudal LC contrast, we take caution in interpreting these associations. Further, the imaging method may be less reliable in the LC’s caudal aspect, as this aspect is more diffusely organized relative to the rostral aspect (Fernandes et al., 2012).

The present study has several additional limitations. For one, we cannot conclude that the patterns we observed for the LC are unique compared to those for other brain regions. However, the pattern of results did not change when accounting for total ventricular volume, which demonstrates non-specific age-related increases, suggesting that LC associations with thickness may extend beyond those of regions which change regularly with age. In addition, we have not included cognitive measures as the BASE-II cognitive battery was designed with a focus on episodic memory rather than tasks associated with the frontoparietal network. However, using a 99% overlapping sample from BASE-II, we recently reported an association between LC contrast and initial recall on the Rey Auditory Verbal Learning Task (Dahl et al., 2019), which may also depend on attention (Chun & Turke-Browne, 2007). Another limitation is that due to the smaller size of the younger relative to the older cohort, results of analyses performed on the entire sample may be driven by older adults. In addition, we cannot rule out the potential influence of external factors such as time of day which may explain some variability in measurements of brain structure (Karch et al., 2019). Finally, both the younger and older adults who participated in BASE-II constituted a highly educated convenience sample (Bertram et al., 2014), meaning that results may not generalize to the general population.

We have offered hypotheses about processes underlying a relationship between LC neuronal integrity and cortical thickness, but these possibilities remain to be tested. In addition, it remains to be understood precisely what is reflected by MRI-based measures of LC contrast. Measures of LC contrast are interpreted as indices of LC neuronal integrity based on findings of Keren et al. (2015) demonstrating that locations of high LC contrast correspond to locations of cells in the LC which contain neuromelanin, a pigment byproduct of catecholamine breakdown (Fedorow et al., 2005; Mann & Yates, 1974). One other possibility is that the LC contrast measure reflects other features of noradrenergic neurons (e.g., water content), besides neuromelanin (Watanabe et al., 2019a; Watanabe et al., 2019b). Furthermore, there has been limited work relating MRI-based LC contrast measures to LC function. One study demonstrated that LC contrast was associated with LC activity during goal-relevant processing (Clewett et al., 2018). Another recent study found that functional connectivity between LC and FPN brain regions during an auditory oddball task was associated with LC contrast (Mather et al., 2020). However, more work is needed to understand the LC contrast-LC function relationship.

To conclude, we found that LC MRI contrast, particularly that of the rostral LC, was associated with thickness of widespread cortical regions in older adults. These findings constitute novel evidence for a link between LC contrast and brain structure in older adulthood and are consistent with previous reports of LC contrast being associated with cognition in older adulthood. Furthermore, our results provide strong motivation for studying the relationship between the LC and FPN in older adulthood, as these brain regions are implicated in attention and memory functions that undergo regular age-related decline.

## Acknowledgements

This article uses data from the Berlin Aging Study (BASE-II) which has been supported by the German Federal Ministry of Education and Research (Bundesministerium für Bildung and Forschung, BMBF) under grant numbers 16SV5537, 16SV5837, 16SV5538, 16SV5536K, 01UW0808, and 01UW070. Additional support was provided by all participating institutions in Germany: Max Planck Institute for Human Development, Humboldt-Universität zu Berlin, Charité-Universitätsmedizin Berlin, Max Planck Institute for Molecular Genetics, Socio-Economic Panel at the German Institute for Economic Research, Universität zu Lübeck, and Universität Tübingen. BASE-II is also part of the BMBF funded EnergI consortium (01GQ1421B). The Responsibility for the contents of this publication lies with its author(s).

This material is based on work supported by the National Science Foundation Graduate Research Fellowship Program under Grant No. DGE-1842487. Any opinions, findings, and conclusions or recommendations expressed in this material are those of the author(s) and do not necessarily reflect the views of the National Science Foundation. SLB’s work was also supported by a Graduate Study Scholarship from the German Academic Exchange Service (DAAD). MJD was a fellow of the Max Planck Research School on the Life Course and received support from the Sonnenfeld-Foundation. MWB’s work was supported by the German Research Foundation (DFG, WE 4269/5-1) and an Early Career Research Fellowship from the Jacobs Foundation. MM’s work was supported by an Alexander von Humboldt fellowship and by National Institutes of Health grant R01AG025340. We thank Michael Krause and Yana Fandakova for assistance with cluster computing, as well as Carsten Finke, Andrei Irimia, and Nils Bodammer for useful discussions and suggestions.

## Data Statement

Data will be shared upon approved request to the BASE-II Steering Committee (https://www.base2.mpg.de/en/project-information/team).

## Abbreviations

BASE-II: Berlin Aging Study-II
LC: locus coeruleus
DPT: dorsal pontine tegmentum
FPN: frontoparietal network
MMSE: Mini Mental State Examination
MRI: magnetic resonance imaging
NE: norepinephrine

