## Supplementary Material for "Locus coeruleus MRI contrast is associated with cortical thickness in older adults"

**Table of Contents**

**1. Supplementary Methods**

**1.1. Visualization of turbo spin echo scans** **3**
*Figure S1*. Turbo spin echo scan from randomly selected older and younger participants

**1.2. Cluster-wise permutation test of age differences in contrast according to LC topography** **3**

**1.3. Analysis of associations between LC contrast and global cortical thickness, controlling for sex** **4**

**1.4. Vertex-wise analysis of associations between LC contrast and cortical thickness, controlling for sex** **5**

**2. Supplementary Results**

**2.1. LC contrast and cortical thickness in the sample** **6**
*Figure S2*. LC contrast versus chronological age in younger and older adults
*Figure S3*. Left and right LC contrast in older and younger adults
*Figure S4*. Global cortical thickness versus chronological age in younger and older adults
*Figure S5*. Overall LC contrast and global cortical thickness by age group and sex

**2.2. Uncorrected significance maps from vertex-wise analysis of associations between LC contrast and cortical thickness** **10**
*Figure S6*. Uncorrected significance maps from analyses of overall LC contrast and thickness
*Figure S7*. Uncorrected significance maps from analyses of rostral LC contrast and thickness
*Figure S8*. Uncorrected significance maps from analyses of caudal LC contrast and thickness

**2.3. Analysis of associations between left and right LC contrast and cortical thickness** **13**
2.3.1. Analysis of left and right LC contrast and global cortical thickness
*Table S1*. Results of regression analyses examining the association between left LC contrast and global cortical thickness
*Table S2*. Results of regression analyses examining the association between right LC contrast and global cortical thickness
2.3.2. Analysis of left and right LC contrast and global cortical thickness
*Table S3*. Cortical clusters where the left LC contrast-thickness association was greater in older than in younger adults (A) and positive in older adults (B)
*Table S4*. Cortical clusters where the right LC contrast-thickness association was greater in older than in younger adults (A) and positive in older adults (B)

**2.4. Analysis of associations between LC contrast and cortical thickness, controlling for sex** **17**
2.4.1. Analysis of LC contrast and global cortical thickness

*Table S5.* Results of regression analyses examining the association between overall LC contrast and global cortical thickness, after controlling for age and sex
*Table S6.* Results of regression analyses examining the association between rostral LC contrast and global cortical thickness, after controlling for age and sex
*Table S7.* Results of regression analyses examining the association between caudal LC contrast and global cortical thickness, after controlling for age and sex
2.4.2. Vertex-wise analysis of LC contrast and regional cortical thickness
*Table S8*. Cortical clusters where the association between overall LC contrast and thickness was greater in older than younger adults (A) and positive in older adults (B)
*Table S9*. Cortical clusters where the association between rostral LC contrast and thickness was greater in older than younger adults (A) and positive in older adults (B) and where caudal LC contrast was associated with thickness in older adults (C).

**<u>3. Supplementary References</u>** **25**

### 1. Supplementary Methods

#### 1.1. Visualization of turbo spin echo scans

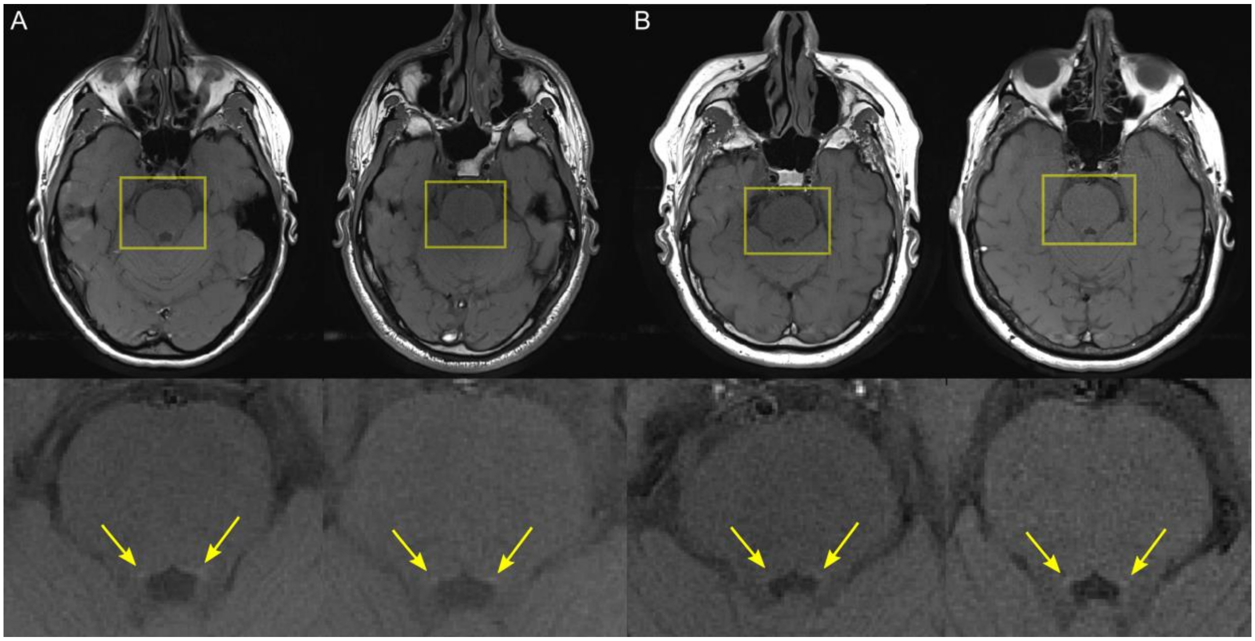


*Figure S1*. Turbo spin echo scans from two randomly selected younger (A) and two randomly selected older (B) adult participants. Axial view of scans is shown in upper panel. Closer view of pontine region in the same scans, outlined in yellow, is shown in lower panel. In lower panel, yellow arrows indicate hyperintensities bordering the fourth ventricle.

#### 1.2. Cluster-wise permutation test of age differences in contrast according to LC topography

To identify clusters of slices where LC contrast differed significantly between younger and older adults, we reimplemented the nonparametric, cluster-wise permutation test also performed in Dahl et al. (2019), using the FieldTrip toolbox (Oostenveld et al., 2011; http://www.fieldtriptoolbox.org). This entailed first performing an independent samples *t*-test comparing LC contrast of older and younger adults for each slice where all participants exhibited reliable LC signal intensity (n=16 slices). Next, clusters were formed by grouping adjacent slices where $p < 0.05$, and a cluster mass statistic for each cluster was calculated by summing the *t*-statistic values of all slices comprising each cluster. A Monte Carlo method was used to generate a permutation distribution of mass statistics for the summed *t*-statistics in each cluster under the assumption of no effect of age group on LC contrast; specifically, in each of 100,000 permutations, age group labels were shuffled, slice-wise independent samples *t*-tests were performed, and cluster mass statistic(s) were computed. To implement a two-tailed test, we considered a cluster significant, if its cluster mass statistic fell in the lowest or highest 2.5% of the respective permutation distribution. For each resulting cluster(s) where LC contrast differed significantly between older and younger adults, we averaged LC contrast values over the slices comprising each cluster to obtain cluster-specific LC contrast estimates for each participant.

**1.3. Analysis of associations between LC contrast and global cortical thickness, controlling for sex**

Exploratory analyses of our data revealed significant sex differences in both LC contrast (Figure S3A) and global cortical thickness (Figure S3B), with females demonstrating significantly greater values of both measures than males. Consequently, in an attempt to test whether the LC contrast-thickness associations depended on sex, we performed a series of supplemental analyses including sex as a predictor of thickness. However, we advise caution in the interpretation of these results, because LC contrast and sex were collinear in both age groups. Having correlated (e.g. non-independent) predictors in a regression analysis can lead to unreliable estimates of the associations between the individual predictors and the response variable (James et al., 2013).

Similar to the approach taken in the main text, we first performed a multiple linear regression analysis using data from all participants to examine the extent to which overall LC contrast and chronological age predicted global thickness and to what extent the overall LC contrast-thickness association differed by age group. We added sex as a predictor, as well as interaction effects between (a) sex and overall LC contrast, (b) sex and age group, and (c) a three-way interaction effect between sex, age group, and overall LC contrast. For these analyses, sex was coded as female = 1, male = -1 so that regression coefficients for overall LC contrast, age, and the overall LC contrast-by-age interaction would reflect values averaged across females and males. As with all other analyses involving age group, age group was coded as older adults = 1, younger adults = -1. We then repeated this analysis separately for rostral and caudal LC contrast values.

**1.4. Analysis of associations between LC contrast and global cortical thickness, controlling for sex**

To examine cortical regions where the association between LC contrast and thickness differed in older and younger adults after accounting for effects of chronological age and sex on thickness, we performed a vertex-wise analysis using Freesurfer's group analysis stream. For this analysis, four groups were defined (older adult females, older adult males, younger adult females, and younger adult males), and the LC contrast slope of the former two groups was contrasted with that of the latter two groups, controlling for effects of chronological age. Consistent with analyses of global thickness, we then examined cortical regions where LC contrast was associated with thickness after accounting for age and sex in each group separately. The procedure was similar to that used in main analyses, but in addition to age being included as a covariate, sex was included as a fixed factor. Cluster-wise correction for multiple comparisons was performed in the same method as described in the main text. These analyses were repeated separately to identify cortical regions where thickness was associated with each rostral and caudal LC contrast.

### 2. Supplementary Results

#### 2.1. LC contrast and cortical thickness in the sample

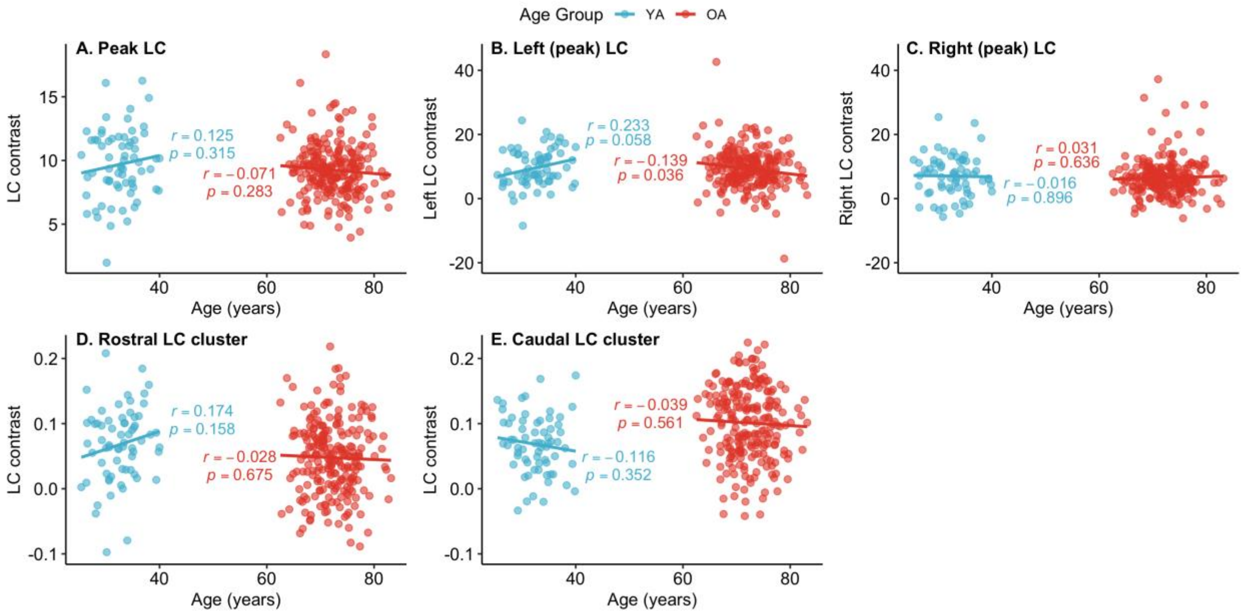


*Figure S2.* Locus coeruleus (LC) contrast versus chronological age in younger adults (YA) and older adults (OA). Pearson's correlation analyses indicated that chronological age was not significantly correlated with overall LC contrast in either younger adults, $r(65) = 0.125$, $p = .315$, or older adults, $r(227) = -0.071$, $p = .283$ (A). Left LC contrast was negatively correlated with age in older adults, $r(227) = -0.139$, $p = .036$, but was not significantly correlated with age in younger adults, $r(65) = 0.233$, $p = .058$ (B). Right LC contrast was not significantly correlated with age in older adults, $r(227) = 0.031$, $p = .636$, or younger adults, $r(65) = -0.016$, $p = .896$ (C). Rostral LC contrast was not significantly correlated with age in younger adults, $r(65) = 0.174$, $p = .158$, or older adults, $r(227) = -0.028$, $p = .675$ (D). Caudal LC contrast was also not significantly correlated with age in younger adults, $r(65) = -0.116$, $p = .352$, or older adults, $r(227) = -0.039$, $p = .561$ (E). Fisher's *r*-to-*z* transformations indicated that the correlation between age and overall LC contrast differed significantly for older and younger adults, $Z = 2.38$, $p = .020$, as did the correlation between rostral LC contrast and age, $Z = 2.47$, $p = .010$, and the correlation between left LC contrast and age, $Z = 4.56$, $p < .001$. The correlation between age and caudal LC contrast did not differ significantly between older and younger adults, $Z = 0.940$, $p = .350$, nor did the correlation between age and right LC contrast, $Z = 0.580$, $p = .560$. While the correlations between LC contrast and age did not reach statistical significance, the trends observed for overall and rostral LC contrast are in line with the results reported by Liu et al., 2019 (Figures 5a,b), wherein LC contrast ratios increased with age in younger adults and decreased with age in older adults.

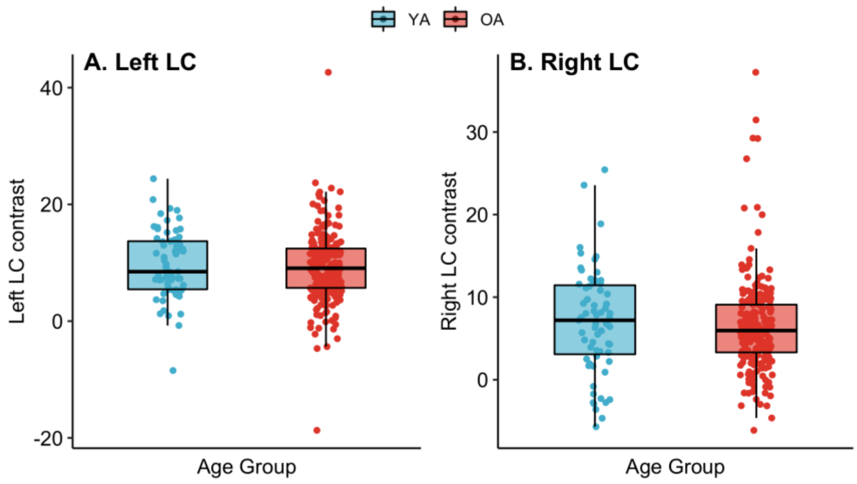


*Figure S3.* Left and right locus coeruleus (LC) contrast in younger adults (YA) and older adults (OA). A 2 (laterality: left, right) x 2 (age group: OA, YA) mixed-design ANOVA indicated a significant main effect of laterality on LC contrast, $F(1, 294) = 45.1$, $p < .001$, with left LC contrast being greater than right LC contrast. There was no significant main effect of age group on LC contrast, $F(1,294) = 0.45$, $p = .503$, and no significant interaction effect between laterality and age group, $F(1,294) = 0.025$, $p = .874$.

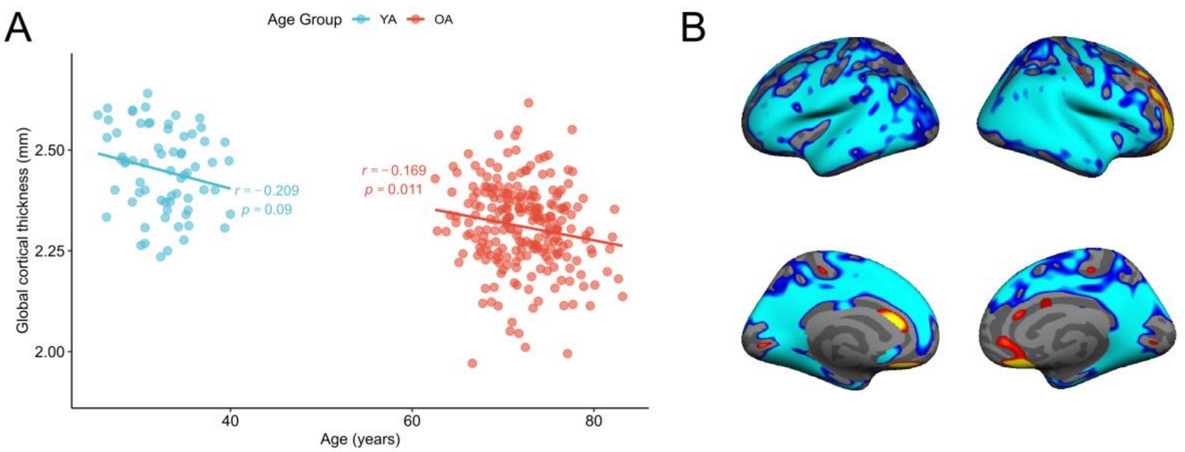


*Figure S4.* (A) Global cortical thickness versus chronological age in younger adults (YA) and older adults (OA). Pearson's correlation analyses demonstrated that chronological age was negatively correlated with global cortical thickness in older adults, $r(227) = -0.169$, $p = .011$. Further, we observed a trend towards a negative correlation between age and thickness in younger adults, $r(65) = -0.209$, $p = .090$. A Fisher's *r*-to-*z* transformation revealed that the correlation between age and thickness did not differ significantly between older and younger adults, $Z = 0.510$, $p = .610$. (B) Uncorrected significance maps (vertex-wise $p < .05$) depicting cortical vertices where thickness differed in older versus younger adults; cooler colors (blue-turquoise) depict vertices where thickness was lower in older than in younger adults, and warmer colors (red-yellow) indicate vertices where thickness was higher in older than in younger adults.

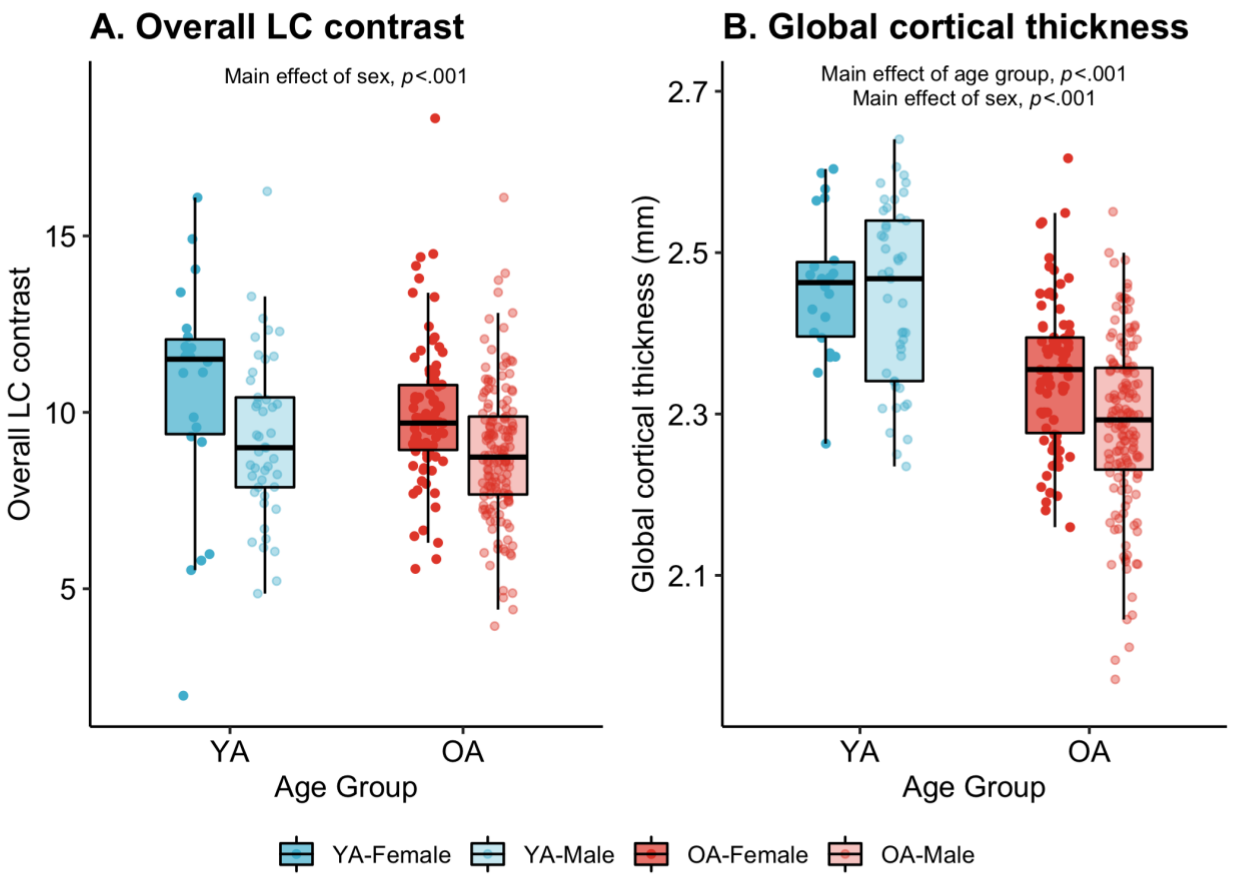


*Figure S5.* Overall locus coeruleus (LC) contrast and global cortical thickness by age group and sex. As an exploratory analysis of whether sex had an effect on overall LC contrast in either age group, we performed a series of 2 (age group: OA, YA) x 2 (sex: female, male) analyses of variance (ANOVAs). (A) The first ANOVA revealed a main effect of sex on overall LC contrast, $F(1, 292) = 18.5$, $p < .001$, with females having greater contrast than males. This analysis indicated no significant main effect of age group, $F(1, 292) = 2.23$, $p = .136$, or interaction effect between age group and sex on overall LC contrast, $F(1, 292) = 0.126$, $p = .723$. (B) The second ANOVA indicated a main effect of sex, $F(1, 292) = 17.8$, $p < .001$, a main effect of age group, $F(1, 292) = 96.5$, $p < .001$, and a trend towards an interaction effect of age group and sex on global thickness, $F(1, 292) = 2.87$, $p = .092$. On boxplots, upper and lower hinges correspond to first and third quartiles, respectively, and medians are indicated with horizontal bars. OA = older adults; YA = younger adults.

**2.2. Uncorrected significance maps from vertex-wise analyses of associations between LC contrast and cortical thickness**

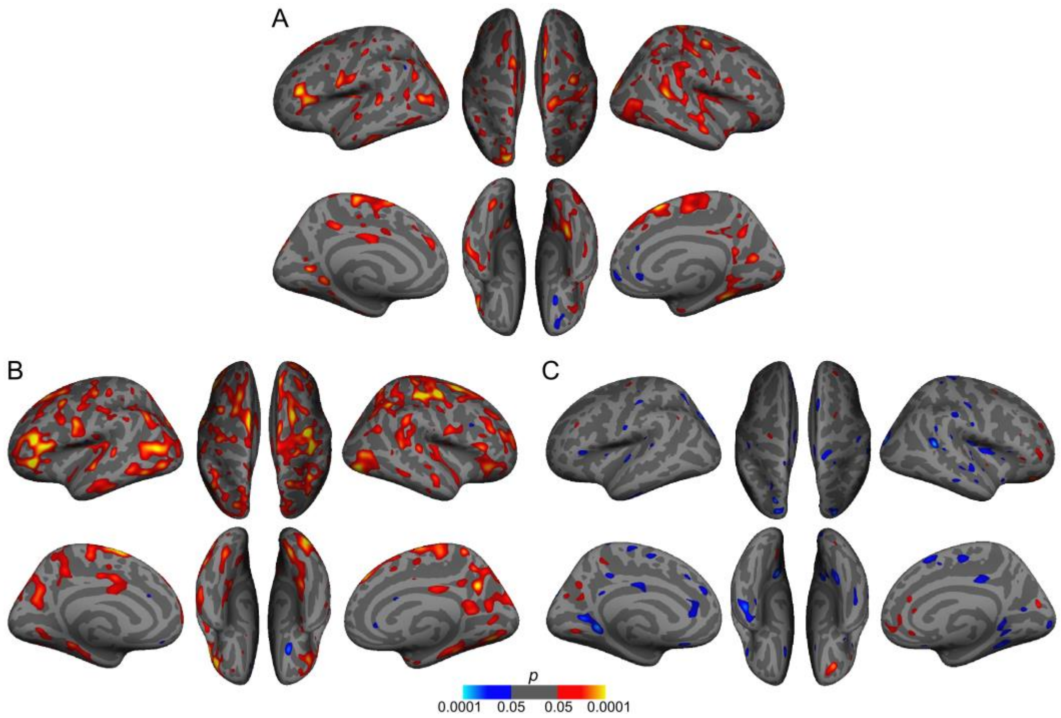


*Figure S6.* Uncorrected significance maps from analyses of overall locus coeruleus (LC) contrast and thickness. (A) shows vertices where the association between overall LC contrast and thickness differed significantly in older and younger adults; in (A), warmer colors (red-yellow) indicate a more positive association in older than younger adults, and cooler colors (blue-turquoise) indicate a more positive association in younger than in older adults. Vertices exhibiting a significant association between overall LC contrast and thickness in older and younger adults are shown in (B) and (C), respectively. In (B) and (C), warmer colors (red-yellow) indicate a positive association between overall LC contrast and cortical thickness, and cooler colors (blue-turquoise) indicate a negative association between overall LC contrast and cortical thickness. All maps were thresholded at vertex-wise $p < .05$.

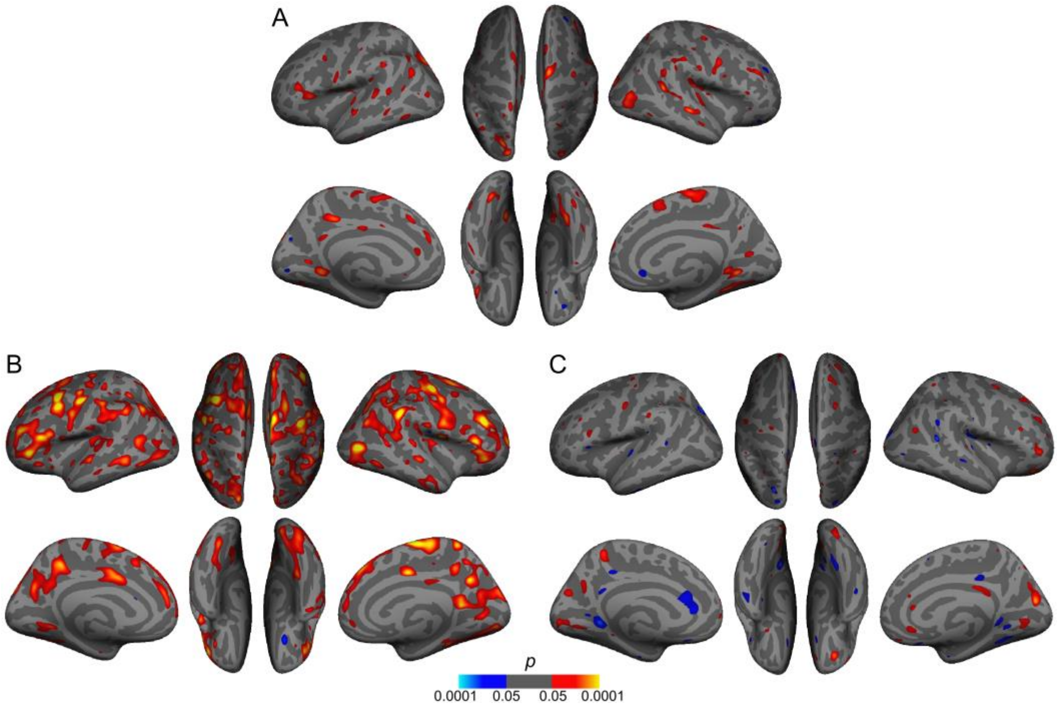


*Figure S7*. Uncorrected significance maps from analyses of rostral locus coeruleus (LC) contrast and thickness. (A) shows vertices where the association between rostral LC contrast and thickness differed significantly in older and younger adults; in (A), warmer colors (red-yellow) indicate a more positive association in older than younger adults, and cooler colors (blue-turquoise) indicate a more positive association in younger than in older adults. Vertices exhibiting a significant association between rostral LC contrast and thickness in older and younger adults are shown in (B) and (C), respectively. In (B) and (C), warmer colors (red-yellow) indicate a positive association between rostral LC contrast and cortical thickness, and cooler colors (blue-turquoise) indicate a negative association between rostral LC contrast and cortical thickness. All maps were thresholded at vertex-wise $p < .05$.

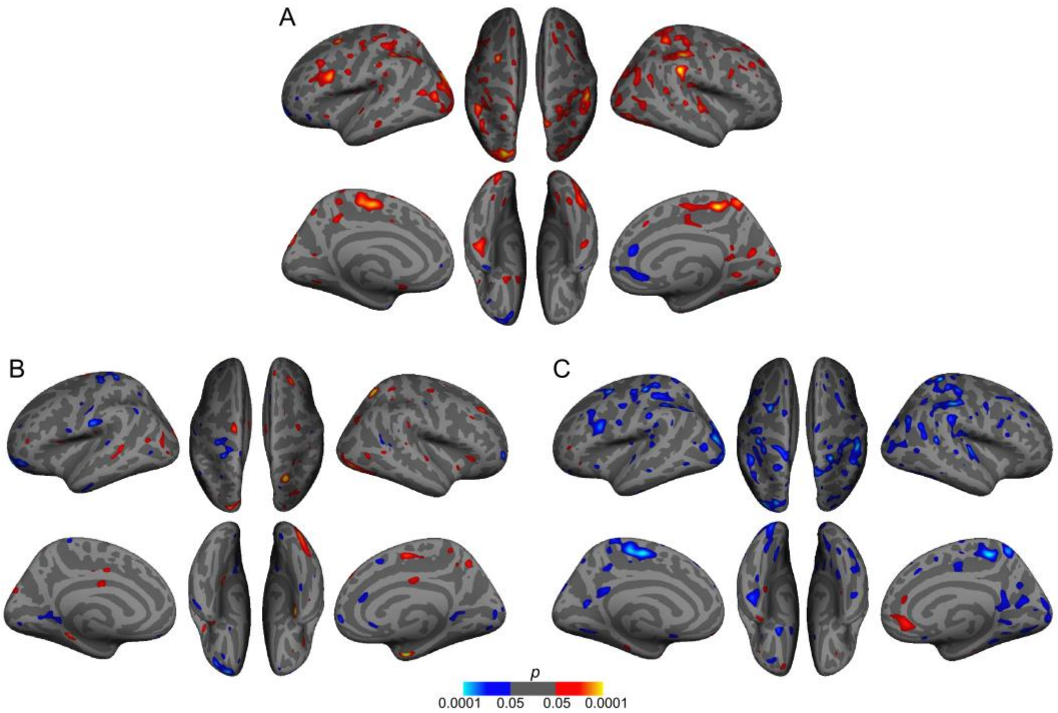


*Figure S8.* Uncorrected significance maps from analyses of caudal locus coeruleus (LC) contrast and thickness. (A) shows vertices where the association between caudal LC contrast and thickness differed significantly in older and younger adults; in (A), warmer colors (red-yellow) indicate a more positive association in older than younger adults, and cooler colors (blue-turquoise) indicate a more positive association in younger than in older adults. Vertices exhibiting a significant association between caudal LC contrast and thickness in older and younger adults are shown in (B) and (C), respectively. In (B) and (C), warmer colors (red-yellow) indicate a positive association between caudal LC contrast and cortical thickness, and cooler colors (blue-turquoise) indicate a negative association between caudal LC contrast and cortical thickness. All maps were thresholded at vertex-wise $p < .05$.

#### 2.3. Analysis of associations between left and right LC contrast and thickness

*2.3.1. Analysis of left and right LC contrast and global cortical thickness*

A series of exploratory multiple regression analyses were performed to examine how each left and right LC contrast were associated with global cortical thickness, after accounting for age. Results of analyses of left and right LC contrast are presented in Tables S1 and S2, respectively. We found a significant interaction effect between left LC contrast and age group on global thickness, indicating the left LC-thickness association differed in older and younger adults (Table S1A). Subsequent analysis in each age group separately indicated that this interaction was driven by a significantly positive association between left LC and global thickness in older adults (Table S1B) but not in younger adults (Table S1C).

An analogous set of tests did not indicate a significant interaction between right LC contrast and age group (Table S2A), although in older adults separately, right LC contrast had a significantly positive association with global thickness (Table S2B). There was no significant association between right LC contrast and global thickness in younger adults (Table S2C). These results indicate that both left and right LC contrast were associated with global cortical thickness in older adults, but a greater positive LC contrast-thickness association in older versus younger adults was only observed for left LC.

Table S1

*Results of multiple linear regression analyses examining the association between left LC contrast and global cortical thickness*

| Predictor | $\beta$ | SE | 95% CI | *t* | *p* |
|---|---|---|---|---|---|
| A. Full sample, $F(3, 292) = 38.98$, $p < .001$, R = 0.286, adjusted $R^2 = 0.279$ | | | | | |
| Left LC contrast | 0.065 | 0.060 | -0.053, 0.183 | 1.09 | 0.278 |
| **Age** | -0.503 | 0.050 | -0.600, -0.406 | -10.2 | <.001 |
| **Left LC contrast * Age group** | 0.120 | 0.060 | 0.002, 0.238 | 2.00 | 0.046 |
| B. Older adults, $F(2, 226) = 8.84$, $p < .001$, $R^2 = 0.073$, adjusted $R^2 = 0.064$ | | | | | |
| **Left LC contrast** | 0.212 | 0.065 | 0.085, 0.340 | 3.28 | 0.001 |
| **Age** | -0.139 | 0.065 | -0.267, -0.012 | -2.15 | 0.033 |
| C. Younger adults, $F(2, 64) = 1.53$, $p = .225$, $R^2 = 0.046$, adjusted $R^2 = 0.016$ | | | | | |
| Left LC contrast | -0.045 | 0.126 | -0.296, 0.206 | -0.36 | 0.723 |
| Age | -0.199 | 0.126 | -0.449, 0.052 | -1.58 | 0.119 |

*Note.* Regression coefficients and standard errors are standardized values. Age group was coded as: older adults = 1, younger adults = -1. SE = standard error.

Table S2

*Results of multiple linear regression analyses examining the association between left LC contrast and global cortical thickness*

| Predictor | *β* | SE | 95% CI | *t* | *p* |
|---|---|---|---|---|---|
| A. Full sample, $F(3, 292) = 36.04$, $p < .001$, $R^2 = 0.270$, adjusted $R^2 = 0.263$ | | | | | |
| Right LC contrast | 0.020 | 0.056 | -0.091, 0.131 | 0.35 | 0.724 |
| **Age** | -0.510 | 0.050 | -0.609, -0.412 | -10.2 | <.001 |
| Right LC contrast * Age group | 0.098 | 0.057 | -0.013, 0.209 | 1.73 | 0.084 |
| B. Older adults, $F(2, 226) = 5.42$, $p = .005$, $R^2 = 0.046$, adjusted $R^2 = 0.037$ | | | | | |
| **Right LC contrast** | 0.132 | 0.065 | 0.004, 0.260 | 2.03 | 0.044 |
| **Age** | -0.173 | 0.065 | -0.301, -0.045 | -2.66 | 0.008 |
| C. Younger adults, $F(2, 64) = 1.81$, $p = .173$, $R^2 = 0.053$, adjusted $R^2 = 0.024$ | | | | | |
| Right LC contrast | -0.099 | 0.122 | -0.342, 0.144 | -0.81 | 0.420 |
| Age | -0.211 | 0.122 | -0.454, 0.032 | -1.73 | 0.088 |

*Note*. Regression coefficients and standard errors are standardized values. Age group was coded as: older adults = 1, younger adults = -1. SE = standard error.

#### *2.3.2. Vertex-wise analysis of left and right LC contrast and regional cortical thickness*

Vertex-wise analyses were performed to determine cortical regions where each left and right LC contrast were associated with thickness, after accounting for the effect of chronological age. Cortical clusters where left and right LC contrast were associated with thickness after cluster-wise multiple comparison correction are presented in Tables S3 and S4, respectively. We identified two clusters where the association between left LC contrast and thickness differed in older versus younger adults (Table S3A); in each of these clusters, the association was more positive in older than younger adults. Further, we identified eleven clusters where left LC contrast was positively associated with thickness in older adults (Table S3B). These clusters corresponded to many of the clusters where overall and rostral LC contrast were associated with thickness in older adults (main text, Tables 5 and 6).

For right LC contrast, the association with thickness differed in older and younger adults in two cortical clusters (Table S4A); in this cluster, the association with right LC contrast was more positive in older versus younger adults. In older adults, right LC contrast was positively associated with thickness in one cortical cluster (Table S4B). There were no clusters where left or right LC contrast was associated with thickness in younger adults. Together, these results indicate that in older adults, left LC was associated with thickness in a more widespread set of cortical regions than was right LC contrast.

Table S3

*Cortical clusters where the association between left LC contrast and thickness was greater in older than in younger adults (A) and positive in older adults (B)*

| Region | Hemi-sphere | Size ($mm^2$) | Peak vertex* | | | CWP** |
|---|---|---|---|---|---|---|
| | | | X | Y | Z | |
| A. Left LC contrast-thickness association greater in OA vs. YA | | | | | | |
| Pars triangularis | Left | 900.35 | -48.7 | 26.7 | 5.4 | 0.0230 |
| Post central | Right | 974.81 | 59.8 | -13.0 | 30.7 | 0.0143 |
| B. Left LC contrast positively associated with thickness in OA | | | | | | |
| Precentral | Left | 5075.26 | -58.5 | 2.9 | 26.2 | 0.0001 |
| Inferior parietal | Left | 4654.32 | -44.2 | -64.5 | 42.0 | 0.0001 |
| Pars triangularis | Left | 3308.43 | -47.9 | 35.6 | -8.6 | 0.0001 |
| Superior parietal | Left | 1044.15 | -23.2 | -84.0 | 25.3 | 0.0098 |
| Precentral | Right | 5166.03 | 40.7 | -10.3 | 59.3 | 0.0001 |
| Inferior parietal | Right | 3241.41 | 35.0 | -74.2 | 40.0 | 0.0001 |
| Rostral middle frontal | Right | 1805.06 | 40.4 | 25.4 | 32.6 | 0.0001 |
| Lateral occipital | Right | 1779.82 | 39.4 | -81.0 | -12.8 | 0.0001 |
| Supramarginal | Right | 1359.55 | 59.9 | -39.9 | 34.5 | 0.0007 |
| Lingual | Right | 1290.46 | 22.3 | -72.4 | -6.2 | 0.0014 |
| Superior parietal | Right | 1135.09 | 23.7 | -85.8 | 30.5 | 0.0051 |

*Note*. OA = older adults; YA = younger adults.
*Talairach coordinates of vertex with peak CWP value.
**Cluster-wise probability (CWP) resulting from cluster-wise Monte Carlo correction for multiple comparisons, reflecting the probability of the cluster appearing by chance. Only clusters with CWP<.05 are listed.

Table S4

*Cortical clusters where the association between right LC contrast and thickness was greater in older than in younger adults (A) and positive in older adults (B)*

| Region | Hemi-sphere | Size ($mm^2$) | Peak vertex* | | | CWP** |
|---|---|---|---|---|---|---|
| | | | X | Y | Z | |
| A. Right LC contrast-thickness association greater in OA vs. YA | | | | | | |
| Fusiform | Right | 1102.77 | 35.6 | -41.9 | -15.5 | 0.0060 |
| Precentral | Right | 994.52 | 40.1 | 3.6 | 13.7 | 0.0128 |
| B. Right LC contrast positively associated with thickness in OA | | | | | | |
| Postcentral | Right | 801.08 | 41.6 | -24.7 | 53.9 | 0.0498 |

*Note*. OA = older adults; YA = younger adults.
*Talairach coordinates of vertex with peak CWP value.
**Cluster-wise probability (CWP) resulting from cluster-wise Monte Carlo correction for multiple comparisons, reflecting the probability of the cluster appearing by chance. Only clusters with CWP < .05 are listed.

### 2.4. Analysis of associations between LC contrast and cortical thickness, controlling for sex

#### *2.4.1. Analysis of LC contrast and global cortical thickness*

Results of the analyses examining the association between overall LC contrast and global cortical thickness including sex as a predictor are presented in Table S5. Overall, these results indicate that accounting for sex did not qualitatively change the associations we observed between overall LC contrast and global thickness. Specifically, after accounting for the effects of sex and its interactions with age and overall LC contrast on global thickness, we found a significant interaction effect between overall LC contrast and age group on global thickness (Table S5A), indicating that the association between overall LC contrast and global thickness differed significantly between older and younger adults. In this analysis, sex and age were also significant predictors of global thickness, but we did not observe a significant interaction effect between overall LC contrast and sex or a significant interaction effect between overall LC contrast, age group, and sex. Thus, we performed post hoc regression analyses separately for each age group, including overall LC contrast, age, and sex as predictors of global thickness. These analyses indicated that overall LC contrast was positively associated with global thickness in older adults (Table S5B), even after accounting for effects of age and sex on global thickness. However, in younger adults, a similar model predicting global thickness from overall LC contrast, age, and sex fit the data poorly, with none of these three variables being significantly associated with global thickness (Table S5C).

Table S6 presents results of similar regression analyses examining the association between rostral LC contrast and global cortical thickness after accounting for age and sex. In this case, after taking sex into account, we did not observe a significant rostral LC contrast-by-age group interaction (Table S6A), which approached significance when sex was not included as a predictor (main text, Table 3). Consistent with analyses for overall LC contrast, we examined the association between rostral LC contrast and global thickness separately in each age group, including age and sex as predictors. These analyses demonstrated that in older adults, after accounting for age and sex effects on thickness, higher rostral LC contrast significantly predicted greater cortical thickness (Table S6B), whereas in younger adults, neither age, sex, nor rostral LC contrast were significant predictors of global thickness (Table S6C). These results indicate that the positive association we observed between rostral LC contrast and global thickness in older adults persisted when also accounting for sex effects on thickness.

Results of analogous regression analyses examining the association between caudal LC contrast and global thickness including age and sex as predictors are presented in Table S7. As in main analyses, we observed a trend towards caudal LC contrast being negatively associated with global thickness; however, did not observe a significant interaction effect between caudal LC contrast and age group on global thickness (Table S7A), whereas this interaction effect approached significance when sex was not included as a predictor (main text, Table 4). Subsequent regression analyses in each age group separately indicated that, as was the case in main analyses, the effect of caudal LC contrast was driven by younger adults, with lower caudal LC contrast in younger adults being a significant predictor of greater global thickness (Table S7C). In contrast, caudal LC contrast was not significantly associated with global thickness in older adults after regressing out effects of age and sex on global thickness (Table S7B). These results indicate that the negative association we observed between caudal LC contrast and global thickness in younger adults persisted when also accounting for sex effects on thickness.

Table S5

*Results of multiple linear regression analyses examining the association between overall LC contrast and global cortical thickness, including sex as a predictor*

| Predictor | *β* | SE | 95% CI | *t* | *p* |
|---|---|---|---|---|---|
| A. Full sample, $F(7, 288)=20.26$, $p<.001$, $R^2=0.33$, adjusted $R^2=0.314$ | | | | | |
| Overall LC contrast | 0.043 | 0.053 | -0.062, 0.147 | 0.80 | .423 |
| **Age** | -0.484 | 0.052 | -0.588, -0.381 | -9.24 | <.001 |
| **Sex** | 0.127 | 0.063 | 0.003, 0.252 | 2.01 | .045 |
| **Overall LC contrast * Age group** | 0.121 | 0.053 | 0.017, 0.225 | 2.29 | .023 |
| Overall LC contrast * Sex | 0.086 | 0.053 | -0.019, 0.190 | 1.62 | .107 |
| Age group * Sex | 0.096 | 0.063 | -0.027, 0.220 | 1.53 | .126 |
| Overall LC contrast * Age group * Sex | -0.016 | 0.053 | -0.120, 0.088 | -0.31 | .760 |
| B. Older adults, $F(3, 225)=11.67$, $p<.001$, $R^2=0.135$, adjusted $R^2=0.123$ | | | | | |
| **Overall LC contrast** | 0.151 | 0.064 | 0.024, 0.278 | 2.35 | .020 |
| **Age** | -0.162 | 0.062 | -0.284, -0.039 | -2.60 | .010 |
| **Sex** | 0.263 | 0.067 | 0.132, 0.395 | 3.94 | <.001 |
| C. Younger adults, $F(3, 63)=1.16$, $p=.332$, $R^2=0.052$, adjusted $R^2=0.007$ | | | | | |
| Overall LC contrast | -0.097 | 0.128 | -0.352, 0.159 | -0.75 | .454 |
| Age | -0.189 | 0.128 | -0.444, 0.067 | -1.47 | .145 |
| Sex | 0.040 | 0.138 | -0.235, 0.315 | 0.29 | .773 |

*Note*. Regression coefficients and standard errors are standardized values. Age group was coded as: older adults = 1, younger adults = -1. Sex was coded as females = 1, males = -1. CI = confidence interval; LC = locus coeruleus; SE = standard error.

Table S6

*Results of multiple linear regression analyses examining the association between rostral LC contrast and global cortical thickness, including sex as a predictor*

| Predictor | *β* | SE | 95% CI | *t* | *p* |
|---|---|---|---|---|---|
| A. Full sample, $F(7, 288)=20.21$, $p<.001$, $R^2=0.329$, adjusted $R^2=0.313$ | | | | | |
| Rostral LC contrast | 0.064 | 0.059 | -0.052, 0.180 | 1.09 | .276 |
| **Age** | -0.472 | 0.053 | -0.577, -0.368 | -8.89 | <.001 |
| Sex | 0.109 | 0.064 | -0.018, 0.235 | 1.69 | .092 |
| Rostral LC contrast * Age group | 0.081 | 0.059 | -0.035, 0.197 | 1.38 | .168 |
| Rostral LC contrast * Sex | 0.090 | 0.059 | -0.027, 0.206 | 1.51 | .131 |
| Age group * Sex | 0.124 | 0.064 | -0.001, 0.250 | 1.95 | .052 |
| Rostral LC contrast * Age group * Sex | -0.074 | 0.059 | -0.190, 0.043 | -1.24 | .215 |
| B. Older adults, $F(3, 225)=11.98$, $p<.001$, $R^2=0.138$, adjusted $R^2=0.126$ | | | | | |
| **Rostral LC contrast** | 0.160 | 0.063 | 0.035, 0.284 | 2.52 | .013 |
| **Age** | -0.168 | 0.062 | -0.290, -0.046 | -2.71 | .007 |
| **Sex** | 0.268 | 0.066 | 0.138, 0.398 | 4.07 | <.001 |
| C. Younger adults, $F(3, 63)=0.96$, $p=.416$, $R^2=0.044$, adjusted $R^2=-0.002$ | | | | | |
| Rostral LC contrast | -0.001 | 0.128 | -0.258, 0.255 | -0.01 | .993 |
| Age | -0.206 | 0.130 | -0.465, 0.053 | -1.59 | .117 |
| Sex | 0.013 | 0.137 | -0.261, 0.287 | 0.09 | .925 |

*Note*. Regression coefficients and standard errors are standardized values. Age group was coded as: older adults = 1, younger adults = -1. Sex was coded as females = 1, males = -1.CI = confidence interval; LC = locus coeruleus; SE = standard error.

Table S7

*Results of multiple linear regression analyses examining the association between caudal LC contrast and global cortical thickness, including sex as a predictor*

| Predictor | *β* | SE | 95% CI | *t* | *p* |
|---|---|---|---|---|---|
| A. Full sample, *F*(7, 288)=19.66, p<.001, $R^2$=0.323, adjusted $R^2$=0.307 | | | | | |
| Caudal LC contrast | -0.151 | 0.085 | -0.319, 0.018 | -1.76 | .079 |
| **Age** | -0.432 | 0.058 | -0.545, -0.319 | -7.51 | <.001 |
| **Sex** | 0.172 | 0.070 | 0.035, 0.310 | 2.47 | .014 |
| Caudal LC contrast * Age group | 0.101 | 0.086 | -0.068, 0.271 | 1.18 | .239 |
| Caudal LC contrast * Sex | -0.029 | 0.086 | -0.199, 0.141 | -0.34 | .735 |
| Age group * Sex | 0.111 | 0.069 | -0.026, 0.247 | 1.59 | .112 |
| Caudal LC contrast * Age group * Sex | -0.046 | 0.087 | -0.217, 0.124 | -0.54 | .592 |
| B. Older adults, *F*(3, 225)=9.69, p<.001, $R^2$=0.114, adjusted $R^2$=0.103 | | | | | |
| Caudal LC contrast | -0.032 | 0.063 | -0.157, 0.093 | -0.50 | .616 |
| **Age** | -0.174 | 0.063 | -0.298, -0.050 | -2.77 | .006 |
| **Sex** | 0.308 | 0.066 | 0.178, 0.437 | 4.67 | <.001 |
| C. Younger adults, *F*(3, 63)=2.83, p=.046, $R^2$=0.119, adjusted $R^2$=0.077 | | | | | |
| **Caudal LC contrast** | -0.276 | 0.119 | -0.514, -0.037 | -2.31 | .024 |
| Age | -0.236 | 0.122 | -0.480, 0.008 | -1.94 | .057 |
| Sex | 0.022 | 0.128 | -0.234, 0.278 | 0.17 | .863 |

*Notes.* Regression coefficients and standard errors are standardized values. Age group was coded as: older adults = 1, younger adults = -1. Sex was coded as females = 1, males = -1. CI = confidence interval; LC = locus coeruleus; SE = standard error.

*2.4.2. Vertex-wise analysis of LC contrast and regional cortical thickness*

Vertex-wise analyses indicated that, after accounting for age and sex and correcting for multiple comparisons, the association between overall LC contrast and thickness was greater for older than younger adults in five cortical clusters. The size and details of these clusters which were located within frontal and parietal cortices are presented in Table S8A. Subsequent analysis of older adults only indicated that overall LC contrast was positively associated with cortical thickness in seven cortical clusters. These clusters were located in regions within frontal, parietal, and occipital cortices, and their location and size details are presented in Table S8B. Compared to results of main analyses in which sex was not considered, we observed similar associations in frontal, parietal, and occipital cortices, although the number of clusters observed was reduced when taking sex into account. In younger adults, we did not observe any cortical clusters where overall LC contrast was associated with cortical thickness after multiple comparison correction. Subsequently we examined where on the cortical surface rostral and caudal LC contrast were associated with thickness after controlling for age and sex. We found that rostral LC contrast was more positively associated with thickness in older than younger adults in one cluster in left superior parietal cortex (Table S9A); when examining older adults only, we identified ten clusters where rostral LC contrast was positively associated with thickness (Table S9B), many of which are similar to the regions observed in main results. In addition, there were 3 clusters where caudal LC contrast was negatively associated with thickness in older adults (Table S9C). After controlling for and sex, there were no cortical clusters where the association between caudal LC contrast and thickness differed in older and younger adults or where caudal LC contrast was associated with thickness in younger adults.

Table S8

*Cortical clusters where the association between overall LC contrast and thickness was more positive in older than in younger adults (A) and positive in older adults (B), after controlling for age and sex*

| Region | Hemi-sphere | Size ($mm^2$) | Peak vertex* | | | CWP** |
|---|---|---|---|---|---|---|
| | | | X | Y | Z | |
| A. Overall LC contrast-thickness association greater in OA vs. YA | | | | | | |
| Superior parietal | Left | 1002.58 | -12.8 | -88.5 | 26.4 | 0.0121 |
| Rostral middle frontal | Left | 922.37 | -35.9 | 30.7 | 14.1 | 0.0208 |
| Banks of superior temporal sulcus | Right | 1320.74 | 45.6 | -43.1 | 13.6 | 0.0015 |
| Fusiform | Right | 806.93 | 36.5 | -42.9 | -13.8 | 0.0453 |
| Transverse temporal | Right | 798.86 | 49.7 | -14.5 | 2.6 | 0.0477 |
| B. Overall LC contrast associated with thickness in OA | | | | | | |
| Rostral middle frontal | Left | 1521.81 | -36.4 | 31.5 | 16.2 | 0.0003 |
| Lateral occipital | Left | 1074.83 | -46.0 | -71.1 | 8.6 | 0.0076 |
| Superior parietal | Left | 970.91 | -15.2 | -73.7 | 45.4 | 0.0153 |
| Precuneus | Right | 1469.78 | 15.3 | -54.6 | 13.1 | 0.0002 |
| Fusiform | Right | 1403.35 | 37.4 | -39.5 | -14.9 | 0.0003 |
| Precentral | Right | 1222.49 | 40.4 | -10.9 | 57.9 | 0.0020 |
| Lateral occipital | Right | 1158.57 | 36.7 | -85.4 | -13.8 | 0.0034 |

*Talairach coordinates of vertex with peak CWP value.
**Cluster-wise probability (CWP) resulting from cluster-wise Monte Carlo correction for multiple comparisons, reflecting the probability of the cluster appearing by chance. Only clusters with CWP<.05 are listed.

Table S9

*Cortical clusters where the association between rostral LC contrast and thickness was greater in older than in younger adults (A) and positive in older adults (B) and where the association between caudal LC contrast and thickness was negative in older adults (C), after controlling for age and sex*

| Region | Hemi-sphere | Association with thickness | Size ($mm^2$) | Peak vertex* | | | CWP** |
|---|---|---|---|---|---|---|---|
| | | | | X | Y | Z | |
| A. Rostral LC-thickness association greater in OA vs. YA | | | | | | | |
| Superior parietal | Left | Positive | 859.78 | -23.1 | -73.0 | 29.4 | 0.0342 |
| B. Rostral LC contrast associated with thickness in OA | | | | | | | |
| Pars opercularis | Left | Positive | 1727.08 | -49.9 | 25.4 | 12.0 | 0.0001 |
| Superior parietal | Left | Positive | 1215.68 | -10.8 | -86.7 | 28.7 | 0.0031 |
| Caudal middle frontal | Left | Positive | 830.72 | -39.4 | 10.8 | 52.0 | 0.0337 |
| Supramarginal | Left | Positive | 830.63 | -47.3 | -25.5 | 33.4 | 0.0337 |
| Supramarginal | Right | Positive | 1559.34 | 57.4 | -40.3 | 34.3 | 0.0003 |
| Precuneus | Right | Positive | 1475.78 | 6.1 | -58.4 | 31.2 | 0.0003 |
| Fusiform | Right | Positive | 1459.01 | 34.9 | -39.9 | -17.5 | 0.0004 |
| Precentral | Right | Positive | 1048.24 | 49.7 | -5.1 | 8.5 | 0.0089 |
| Pars triangularis | Right | Positive | 855.06 | 54.0 | 27.0 | 8.3 | 0.0325 |
| Precentral | Right | Positive | 803.61 | 40.8 | -10.3 | 58.2 | 0.0461 |
| B. Caudal LC contrast associated with thickness in OA | | | | | | | |
| Pars orbitalis | Left | Negative | 2298.92 | -36.3 | 48.5 | -9.0 | 0.0001 |
| Postcentral | Left | Negative | 857.06 | -23.6 | -35.9 | 59.9 | 0.0338 |
| Supramarginal | Left | Negative | 835.16 | -58.4 | -21.1 | 25.0 | 0.0373 |

*Talairach coordinates of vertex with peak CWP value.
**Cluster-wise probability (CWP) resulting from cluster-wise Monte Carlo correction for multiple comparisons, reflecting the probability of the cluster appearing by chance. Only clusters with CWP<.05 are listed.
